# Characterization of iGABASnFR2 for in vivo mesoscale imaging of intracortical GABA dynamics

**DOI:** 10.1101/2025.04.22.649661

**Authors:** Edris Rezaei, Setare Tohidi, Mojtaba Nazari-Ahangarkolaee, Javad Karimi Abadchi

**Author notes:** Corresponding and first author: Edris Rezaei, Address: Edris Rezaei, Department of Neuroscience, University of Lethbridge, 4401 University Dr W, Lethbridge, AB T1K 3M4, Canada.

## Abstract

While genetically encoded sensors have advanced the study of cortical excitation, tools for large-scale imaging of inhibition remain limited. Visualizing extracellular GABA dynamics in vivo is essential for understanding how inhibitory networks shape brain activity across sensory, behavioral, and pharmacological states. To validate and apply the genetically encoded sensor iGABASnFR2 for wide-field imaging of extracellular GABA, and to characterize how cortical inhibition reorganizes across brain states, sensory modalities, and after GABA transporter blockade. We performed mesoscale imaging in head-fixed C57BL/6 mice systemically expressing iGABASnFR2. Recordings were conducted under isoflurane anesthesia, during quiet wakefulness, natural sleep (NREM and REM), and after administration of the GAT-1 inhibitor Tiagabine. We analyzed both sensory-evoked and spontaneous GABA signals using time-series, spectral, and seed-pixel correlation analyses. iGABASnFR2 demonstrated strong and modality-specific GABAergic responses to sensory stimulation, with faster and stronger activation in the contralateral cortex. Although the general spatial patterns of sensory-evoked GABA responses were consistent across anesthesia and quiet wakefulness, the amplitude, timing, and spread of these responses were significantly greater during wakefulness. During spontaneous activity, cortical GABA levels and connectivity modulated by brain state: GABA amplitude and interhemispheric synchrony were highest during quiet wakefulness but reduced during NREM sleep. Tiagabine elevated baseline GABA levels, abolished stimulus-evoked responses, and enhanced local and long-range inhibitory synchrony. iGABASnFR2 enables reliable, high-resolution imaging of cortical GABA dynamics in vivo. These results demonstrate that inhibitory signaling is dynamically structured across brain states and can be pharmacologically modulated. This tool offers new opportunities to explore the role of inhibition in health and disease at the mesoscale level.

## Introduction

The nervous system is composed of two basic cell types: neurons and glial cells. Neurons can be categorized into excitatory and inhibitory types. Excitatory neurons primarily release glutamate which is the major neurotransmitter in the nervous system to facilitate communication between neurons across different brain regions^1^, while inhibitory neurons mainly release gamma-aminobutyric acid (GABA) to stabilize neural networks by balancing excitatory activity and preventing excessive neuronal firing^2^. Maintaining this precise balance between excitation and inhibition is essential for sensory processing, memory formation, and motor control^3–4^. Through temporally and spatially precise modulation of neuronal activity, inhibitory signaling contributes to a wide range of brain functions, including sensory processing, circuit refinement, and the regulation of oscillatory dynamics^5–9^. Disruptions in GABAergic signaling underlie a variety of neurological and psychiatric disorders, including epilepsy, schizophrenia, and autism spectrum disorders, highlighting the clinical importance of understanding GABAergic modulation in both health and disease^10–16^. Despite the importance of GABA in regulating cortical activity, direct, real-time visualization of GABA dynamics in vivo remains unknown. The development of genetically encoded neurotransmitter sensors has significantly expanded the ability to monitor neural activity across spatial and temporal scales^17^. For instance, the glutamate sensor iGluSnFR enabled high-resolution, real-time imaging of excitatory neurons, providing insights into cortical connectivity and sensory-evoked activity in awake and anesthetized animals^18–19^. Building on advances in sensor engineering, the genetically encoded fluorescent sensor iGABASnFR2 was developed to detect extracellular GABA dynamics^20^. This sensor permits real-time, in vivo monitoring of extracellular GABA, providing a powerful tool for visualizing GABA’s spatial and temporal dynamics across large-scale cortical regions. Wide field imaging with iGABASnFR2 enables comprehensive mapping of inhibitory circuits. However, a detailed characterization of iGABASnFR2 functionality across different brain states, sensory modalities, and pharmacological conditions is still needed. In this study, we comprehensively characterized spontaneous and sensory-evoked GABAergic activity across the dorsal cortex using wide-field imaging of iGABASnFR2 sensor in natural sleep, awake mice, and under 1% isoflurane anesthesia. Additionally, we assess the effects of Tiagabine, a GABA reuptake inhibitor, to further characterize sensor performance and demonstrate its sensitivity to pharmacological manipulation of cortical GABA levels. These results not only validate the robustness and sensitivity of iGABASnFR2 as a practical tool for monitoring GABAergic neurotransmission in vivo but also lay a solid foundation for future studies into inhibitory dynamics in health and disease, including conditions like epilepsy, schizophrenia, and autism spectrum disorders.

## Material and methods

### Animal subjects

The University of Lethbridge animal care committee approved all procedures, which adhered to the guidelines set forth by the Canadian Council on Animal Care and Use. We used 20 young adult (6–8 weeks old) C57BL/6 mice (12 males and 8 females). Mice were accommodated in transparent plastic cages within a 12-hour light-dark cycle, with lights turning on at 7:30 AM and provided unrestricted food and water access. Room temperature was maintained at 24 ± 2 °C, and relative humidity was kept between 40% and 50%.

### Viral constructs

The viral constructs originated from the Viral Vector Core of the Canadian Neurophotonics Platform (RRID: SCR_016477). Plasmids encoding iGABASnFR2 and csGFP were obtained from Addgene, and AAV2/PHP.N-CAG-iGABASnFR2 and AAV2/PHP.N-CAG-csGFP viral vectors were subsequently designed, packaged, and purified by the CNP Viral Vector Core at a final concentration of 1.5 × 10¹³ GC/mL.

### Retro-orbital injection

For retro-orbital viral injections^21^, 4–6-week-old C57BL/6 mice were anesthetized with 3% isoflurane and maintained at 2–2.5% during the procedure. Metacam (5 mg/kg, subcutaneous) was administered for analgesia, and body temperature was maintained using a heating blanket. Topical anesthesia (0.5% proparacaine hydrochloride) was applied to the eye, followed by gentle pressure to induce mild eye protrusion. A 30-gauge needle was then inserted through the medial canthus at a 30° angle into the retro-orbital sinus. Mice received 1.4 × 10¹¹ genome copies (GC) of either AAV2/PHP.N-CAG-iGABASnFR2 or AAV2/PHP.N-CAG-csGFP, using the PHP.N capsid for widespread CNS expression, as originally described by^22^.

### Drug Administration

Tiagabine hydrochloride (Cat #SML0035, Sigma-Aldrich Canada) was purchased from Sigma-Aldrich. Tiagabine was dissolved in sterile saline to achieve a concentration of 5 mg/ml. For intraperitoneal (IP) injection, mice received Tiagabine at a dosage of 10 mg/kg body weight. Control animals were administered an equivalent volume of sterile saline.

### Surgical procedure and post-operative Care

C57BL/6 mice received buprenorphine (0.05–0.1 mg/kg, subcutaneously) approximately 30 minutes before surgery, followed by anesthesia with isoflurane (1–2% in oxygen) delivered via a nose cone. After shaving and sterilizing the scalp, lidocaine (0.5%, 5 mg/mL, subcutaneously) was administered at the incision site for local anesthesia (0.04–0.08 mL for mice weighing 25–55 g). A midline incision was made to expose the skull, and the overlying skin was carefully removed to avoid damaging the underlying bone. A custom-designed head plate was affixed to the skull using C&B Metabond Quick Base (Parkell, NY, USA) mixed with C&B Metabond Clear L-Powder (3 g, Product Code S399, Parkell, Japan). A sterile 12-mm circular glass coverslip (Carolina Biological Supply, Cat. No. 633005) was placed on the skull surface and sealed in place with the same adhesive. For hippocampal recordings, a bipolar electrode made of two twisted 50 μm Teflon-coated stainless-steel wires (A-M Systems) was slowly inserted through a craniotomy at a 57° angle relative to vertical. Electrode placement was guided by continuous monitoring of signal quality using both visual and auditory feedback. After, the electrode was secured to the skull using Krazy Glue, followed by dental cement. An electromyography (EMG) electrode was also implanted into the neck muscles to monitor muscle activity. Following surgery, animals were housed individually in temperature-controlled recovery cages and received subcutaneous injections of Baytril (enrofloxacin, 10 mg/kg), meloxicam (5 mg/kg), and 1 mL of warm sterile saline. These injections were administered once every 24 hours for three days postoperatively, in accordance with institutional guidelines. After this recovery period, animals were monitored twice daily for the remainder of the experiment.

### iGABASnFR2 imaging under anesthesia

Following, the animals were anesthetized with isoflurane (2.5% for induction, followed by 1% for maintenance). The depth of anesthesia was confirmed by assessing reflexes. Once adequately anesthetized, each mouse was positioned in a head-stage, and the head was securely head-fixed. A homoeothermic blanket was utilized to maintain their body temperature, and isoflurane was administered via a nosepiece. The isoflurane concentration was adjusted to 1% to initiate the imaging procedure. Images were acquired using a microscope consisting of a front-to-front pair of video lenses with a field of view measuring 8.6 × 8.6 mm. The camera’s focal plane was positioned 0.5–1 mm (about 0.04) below the cortical surface. A 12-bit charge-coupled device (CCD) camera (1M60 Pantera Dalsa, Waterloo, ON) and an EPIX E8 frame grabber with XCAP 3.8 imaging software (EPIX, Inc., Buffalo Grove, IL) were used to capture images at a frame rate of 80 Hz. These imaging parameters have been employed in previous studies^23–25^. Carefully designed data collection protocols support the robustness of our findings. Sequential illumination was achieved using alternating blue and green LEDs^26^. The timing of LED alternation is illustrated in Supplementary Figure 1. Blue light (473 nm, filtered through a 467–499 nm bandpass) was used to excite the iGABASnFR2 indicator, and green light (530 nm, filtered through a 527/42 nm bandpass) was used for intrinsic signal imaging of blood volume. A bandpass emission filter (shown in Figure 1) was positioned in front of the CCD camera to enable selective detection of either fluorescence or reflectance signals. Blue and green LEDs were synchronized and alternated on a frame-by-frame basis using TTL triggering, resulting in interleaved acquisition of fluorescence and reflectance images at 40 Hz per channel. Additionally, images of reflectance, crucial for blood artifact corrections, were evaluated within the current pipeline^27–29^. Anesthetized iGABASnFR2 imaging of spontaneous activity was conducted without sensory stimulation for 15-minute sessions.

**Figure 1.**
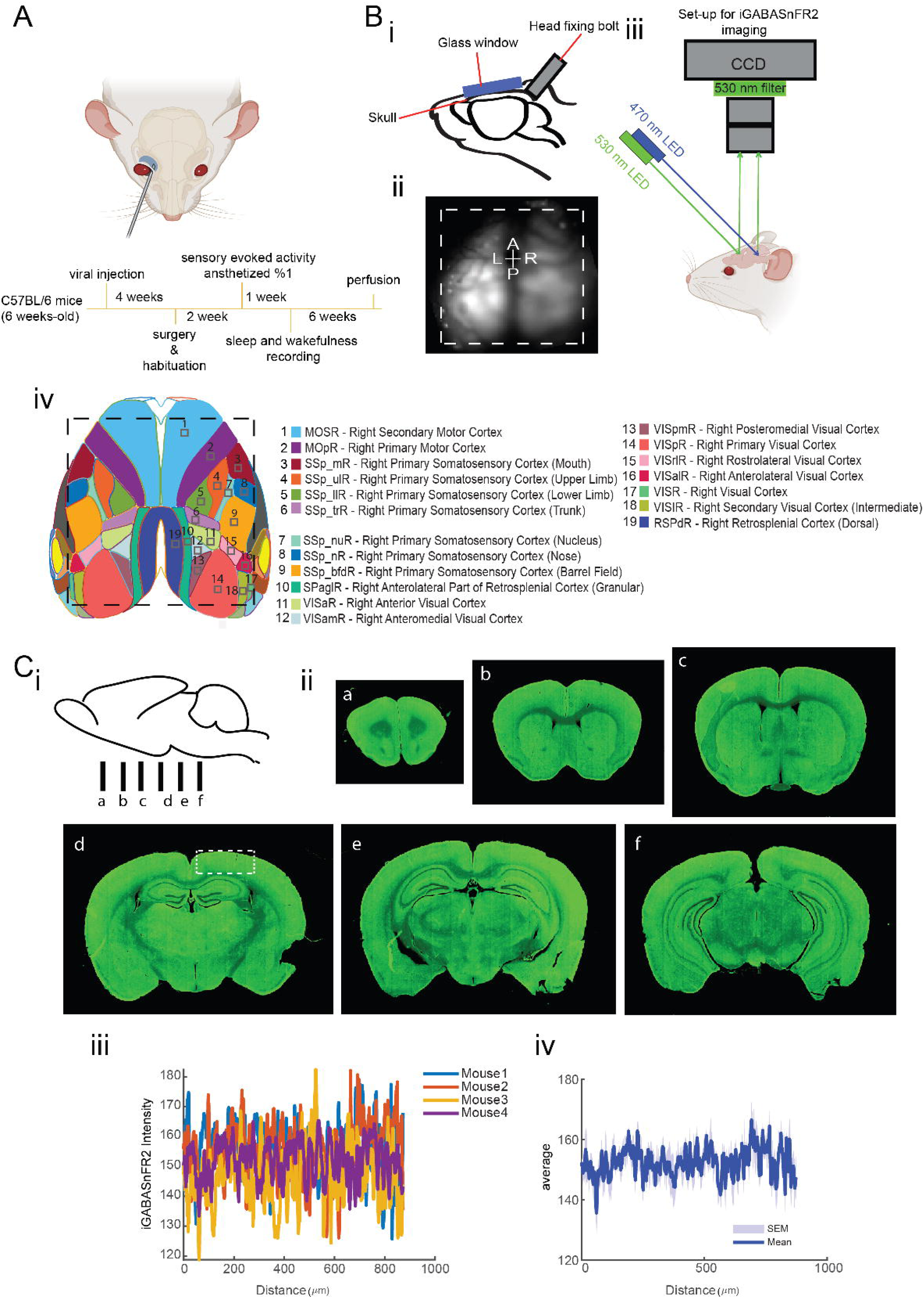
Schematic of experimental workflow, imaging setup, and an expression of iGABASnFR2. A. Experimental timeline. AAV2/PHP.N-CAG-iGABASnFR2 or the control virus AAV2/PHP.N-CAG-csGFP was systemically administered via retro-orbital injection into 6-week-old C57BL/6 mice. Four weeks after viral injection, the animals underwent cranial window surgery. Following a 7-day recovery period, animals gradually habituated to the head-fixation setup over the course of another week. Sensory-evoked imaging was then conducted under 1% isoflurane anesthesia. After this session, longitudinal recordings during quiet wakefulness and natural sleep were carried out for up to six weeks. Mice were subsequently euthanized, and brains were perfused for histological analysis. B. imaging setup. (i) Illustration of the cranial window implantation over the skull following scalp removal. (ii) representative image showing the cortical imaging area, indicated by the dashed white line. (iii) Schematic of the wide-field imaging setup: a blue LED (470 nm) was used for excitation, and signals were captured using a CCD camera at 530 nm emission. (iv) Map of the bilateral craniotomy showing targeted cortical regions based on the Allen Mouse Brain Atlas reference. C. expression of iGABASnFR2. (i) Schematic of coronal sectioning locations (a–f) along the anterior-posterior axis. (ii) Coronal sections (a–f) show strong iGABASnFR2 expression in the cortex and hippocampus. (iii) The region marked by a white dashed rectangle in panel C(iid) was used to extract fluorescence profiles across animals (n=4), highlighting unique expression and inter-animal expression consistency. (iv) The average profile with SEM as a shaded region.

### Sensory stimulation

We captured the iGABASnFR2 signal in reaction to varied peripheral simulations while utilizing urethane anesthesia, following the methodology outlined in previous studies. Sensory stimuli were employed to map the functional areas of the hind-limb somatosensory, forelimb somatosensory, auditory, visual, and barrel cortices^23,30^. Sensory stimuli were applied to map cortical regions corresponding to forelimb, hindlimb, whisker, visual, and auditory modalities. For forelimb and hindlimb stimulation, a piezoelectric bending actuator delivered a single 300-ms tap via a square pulse directly to the skin of one forelimb or hindlimb. Whisker stimulation targeted the C2 whisker, which was attached to a piezoelectric actuator (Q220-A4-203YB, Piezo Systems, Inc., Woburn, MA) and deflected using a single 300-ms square pulse. Visual stimuli consisted of a single 20-ms pulse of 435 nm light (LED), delivered at a fixed distance and height relative to the right eye. For each sensory modality, 40 stimulus presentations were delivered with a 10-second interstimulus interval to calculate average cortical responses. The timing of stimulus delivery is illustrated in Supplementary Figure 1.

### Habituation

After the 7-day recovery period from surgery, mice gradually habituated to head restraint in the recording environment. Initially, each mouse was placed individually on the recording platform along with Cheerios cereal, allowing them to explore freely and become comfortable. Mice were progressively acclimated to eating the cereal while head-fixed, beginning with 5-minute sessions and increasing by 5 minutes each day until reaching 60 minutes.

### iGABASnFR2 imaging during wakefulness

Following habituation, wakefulness recordings began. Each animal underwent recording sessions every three days, completing three to four sessions per mouse. Recordings were consistently performed at the same time of day to reduce variability and stress. Between sessions, mice were returned to their home cages for rest and recovery before the next recording. After finishing all wakefulness sessions, mice proceeded to the sleep recording phase.

### iGABASnFR2 imaging during sleep

To optimize conditions for natural sleep under head restraint, mice were transferred from their colony housing to a separate room at noon the day before recording. Sleep was restricted for six hours by gentle stimulation (using a cotton-tip stick) whenever signs of drowsiness were observed. Following 6 hours of sleep deprivation, mice were placed overnight in large, enriched cages containing a running wheel, Cheerios, and a water container to promote exploration and natural sleep. The next morning (∼9:00 AM), animals were transferred to the imaging platform for sleep recordings. Afterward, they were returned to their home cages for at least three days of recovery before any further recordings. This sleep deprivation protocol is commonly used to induce moderate but physiologically meaningful sleep pressure^31^, which is known to trigger a homeostatic increase in slow-wave activity (SWA) during subsequent NREM sleep.

### Preprocessing

Image stacks were first de-interleaved to separate the GABA-sensitive fluorescence signal (blue channel) and the hemodynamic reflectance (green channel) signal. The correct channel assignment was verified by computing pixel-wise correlations between the first frame of each stack and reference images corresponding to each illumination wavelength. To quantify extracellular GABA dynamics, the relative fluorescence change (ΔF/F) was calculated on a per-pixel basis using a 2-second pre-stimulus window to define baseline fluorescence (F). The ΔF/F signal was defined as:

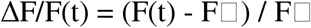

where F(t) is the fluorescence intensity at time t, and F is the baseline fluorescence. To correct for hemodynamic artifacts^27–29^, a pixel-wise linear regression was applied, in which a scaled version of the reflectance signal R(t) was subtracted from the fluorescence trace:

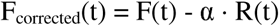

The corrected fluorescence signal was then normalized to the baseline (ΔF/F) and bandpass filtered between 31–74 Hz using a second-order Butterworth filter to remove low-frequency drift and high-frequency noise. Trial-averaged ΔF/F responses were generated to assess sensory-evoked activity. All preprocessed data—including corrected ΔF/F, raw fluorescence, and reflectance signals—were saved in float32 format.

### ROI-based fluorescence analysis

Following preprocessing, imaging data from specific regions of interest (ROIs)—defined as 3×3-pixel areas (∼40,401 µm²) centered around anatomical coordinates corresponding to stimulation sites)— were extracted. Baseline correction was conducted by subtracting the mean fluorescence signals calculated from a 1-second pre-stimulus period from the post-stimulus fluorescence signals. The signals were further filtered to eliminate slow baseline drifts using a high-pass filter (>0.1 Hz) and to reduce high-frequency noise using a low-pass filter (<5 Hz). From the filtered signals, several key parameters were derived, including peak amplitude (maximum ΔF/F within a 1-second post-stimulus interval), time-to-peak, and decay time (duration for the fluorescence to fall to 50% of peak amplitude). For visualization, the corrected ΔF/F signals were plotted with indicators marking peak and decay times, facilitating clear interpretation of response dynamics. Mean responses across trials were computed separately for each sensory modality, with variability assessed by plotting the standard error of the mean (SEM) as shaded regions surrounding the mean trace.

### Motion detection and exclusion

Motion artifacts were identified and excluded from analyses using electromyography (EMG) signals recorded simultaneously with imaging data. EMG recordings were smoothed using a median filter and squared to enhance the detection of muscle activity periods. An activity threshold was established at the 95th percentile of the processed EMG signal to identify movement onset and offset events. These EMG-detected motion periods were temporally aligned with imaging frames via synchronized camera clock signals. Frames coinciding with detected movements were subsequently removed from analysis, ensuring the seed pixel correlation analysis reflected only stationary periods free of motion-related artifacts.

### Seed-pixel correlation analysis

Seed pixel correlation analysis was performed to evaluate functional connectivity based on spontaneous GABA activity from sleep and anesthetized mice. preprocessed data were spatially registered to the Allen Brain Atlas, enabling anatomical alignment and inter-subject comparisons. Regions of interest (ROIs)—specifically the barrel cortex (BC), visual cortex (VC), hindlimb (HL), and forelimb (FL)—were defined using anatomical coordinates derived from the atlas. Within each ROI, seed pixels were selected to serve as reference points for correlation-based connectivity analysis.

### Motion signal extraction, alignment, and sleep scoring

Behavioral videos were used to monitor animal movement during imaging. Motion signals were extracted using FaceMap^32^, which computes frame-to-frame pixel intensity changes in user-defined regions of interest (ROIs). Five ROIs—nose, whisker pad, ear, shoulder, and trunk— were selected to capture both facial and body movements. To synchronize video with neural data, camera frame pulses were recorded on analog channels during acquisition. These were used to align motion traces with electrophysiological recordings (LFP and EMG, sampled at 2 kHz). Traces were interpolated or trimmed as needed, z-scored, and averaged across ROIs to generate composite facial and body motion signals. This enabled detection of both gross movement and small twitches, such as those occurring during REM sleep, which may not be captured by EMG alone. With motion aligned, vigilance states were classified as wakefulness, NREM, or REM sleep using combined behavioral and physiological features. Wakefulness was marked by visible movement and high EMG activity. NREM sleep was defined by low EMG, a low hippocampal theta-to-delta power ratio, and the presence of Large Irregular Activity (LIA) in the LFP. REM sleep was characterized by minimal EMG activity, a high theta-to-delta ratio, and continuous hippocampal theta. In head-fixed recordings, pupil constriction served as an additional marker of sleep onset^33–35^. This multimodal approach enabled robust, accurate classification of sleep stages across different experimental conditions.

### Statistical analysis

All data processing and analyses were performed using custom scripts written in MATLAB R2024a.

## Results

### Experimental workflow, imaging setup, and cortical expression of iGABASnFR2

To characterize extracellular GABA dynamics, we first injected AAV2/PHP.N-CAG-iGABASnFR2 and AAV2/PHP.N-CAG-csGFP systemically via retro-orbital injection (**Figure 1A**). Imaging was performed under isoflurane anesthesia, during quiet wakefulness, NREM and REM sleep. The imaging setup (**Figure 1B**) utilized a bilateral cranial window and a CCD-based wide-field microscope that captured fluorescence across an 8.6 × 8.6 mm field of view. Sequential blue (470 nm) and green (530 nm) LED illumination enabled alternating frame acquisition and hemodynamic correction^27–29^. To validate sensor expression, we conducted a histological analysis (**Figure 1C**). A robust expression of iGABASnFR2 was revealed in cortical and hippocampal regions. Coronal brain sections (**Figure 1Cii**) demonstrated consistent expression across animals. To assess inter-animal variability and regional expression strength, fluorescence intensity profiles were extracted using ImageJ from defined cortical areas (**Figure 1Ciii**), and to further quantify and visualize overall trends, these profiles were averaged across animals, with the mean ± SEM shown (**Figure 1Civ**) confirming uniform sensor expression suitable for quantitative cortical imaging.

### iGABASnFR2 reveals modality– and hemisphere-specific cortical inhibition under anesthesia

Previous studies on sensory processing have primarily focused on excitatory neuronal responses, often using calcium or glutamate indicators to map stimulus-evoked activity across the cortex ^19,23,36^. However, much less is known about how sensory stimuli engage inhibitory networks, particularly at the mesoscale. Inhibitory interneurons play a crucial role in shaping sensory responses, modulating cortical excitability, and controlling the timing and precision of neural coding^37–38^. To examine the spatiotemporal dynamics of GABA activity across the cortex, we delivered contralateral whisker, hindlimb, forelimb, and visual stimulation under 1% isoflurane anesthesia and recorded GABA activity using the iGABASnFR2 sensor. As shown in Figure 2A, each sensory modality shows a localized increase in GABAergic fluorescence within the corresponding primary sensory cortex, with response onsets occurring approximately 75–150 ms after stimulus. All sensory modalities elicited more robust and spatially localized GABAergic activity. Temporal response profiles (**Figure 2B**) demonstrated stronger GABAergic activation in the contralateral hemisphere relative to the ipsilateral side across all sensory modalities. Specifically, the contralateral visual cortex exhibited the highest peak amplitude, followed by hindlimb, whisker, and forelimb cortices while ipsilateral responses were generally weaker and slower. To quantify these differences, we extracted response features including decay time, time to peak, and peak amplitude across all sensory regions (**Figure 2C**). A two-way ANOVA revealed significant main effects of sensory region on all three measures of inhibitory response dynamics. Time to peak (*p* = 0.0092), decay time (*p* = 0.0006), and peak amplitude (*p* = 0.0011) all varied significantly across sensory regions, indicating region-specific characteristics of GABAergic inhibition. Laterality (ipsilateral vs. contralateral) had a significant main effect only on peak amplitude (*p* < 0.0001), with contralateral responses consistently showing higher amplitudes. No significant effects of laterality were observed for time to peak (*p* = 0.35) or decay time (*p* = 0.77). Furthermore, there were no significant interactions between region and laterality for any of the three metrics (time to peak: *p* = 0.22; decay: *p* = 0.83; peak amplitude: *p* = 0.78), suggesting that hemispheric differences in GABAergic inhibition were consistent across modalities and not dependent on specific sensory regions. To confirm that these delayed signals were specific to GABAergic dynamics and not artifacts of hemodynamics or sensor excitation, we used csGFP-expressing mice under identical imaging conditions as a negative control. As shown in Supplementary Figure 2, csGFP mice showed no significant sensory-evoked or spontaneous fluorescence changes, confirming that the iGABASnFR2 signals reflect GABAergic activity. Auditory stimulation under anesthesia evoked clear iGABASnFR2 responses in the auditory cortex (Supplementary Figure 3), further validating sensor specificity. To assess cortical coordination during sensory processing, we performed seed-pixel correlation analysis across 10 anatomically defined cortical regions. This revealed structured and modality-specific inhibitory networks, with the strongest intrahemispheric connectivity observed contralateral to the stimulus (Supplementary Figure 4). Across sensory modalities—including whisker **(Video 1)**, hindlimb (**Video 2)**, forelimb **(Video 3)**, and visual **(Video 4)** stimulation—sensory input activates thalamocortical projections targeting layer 4 of the primary sensory cortices, leading to early excitatory responses^39–40^. These are primarily mediated by excitatory neurons and refined by fast feedforward inhibition from parvalbumin (PV)-expressing interneurons^41–42^. Subsequently, a delayed GABAergic response emerges, largely driven by somatostatin (SST)-expressing interneurons providing feedback inhibition to modulate dendritic activity and maintain cortical stability^9,43,44^. This response is consistently stronger and earlier in the contralateral hemisphere, which receives direct thalamic input, whereas the ipsilateral hemisphere exhibits a weaker and more delayed GABAergic response, likely due to slower callosal transmission and reduced excitatory drive^45–46^. Together, these results demonstrate that sensory-evoked GABA dynamics follow a consistent temporal structure across modalities and hemispheres and confirm that iGABASnFR2 reliably detects extracellular GABA responses with spatiotemporal precision across cortical regions, establishing its utility for mesoscale mapping of inhibitory dynamics.

**Figure 2.**
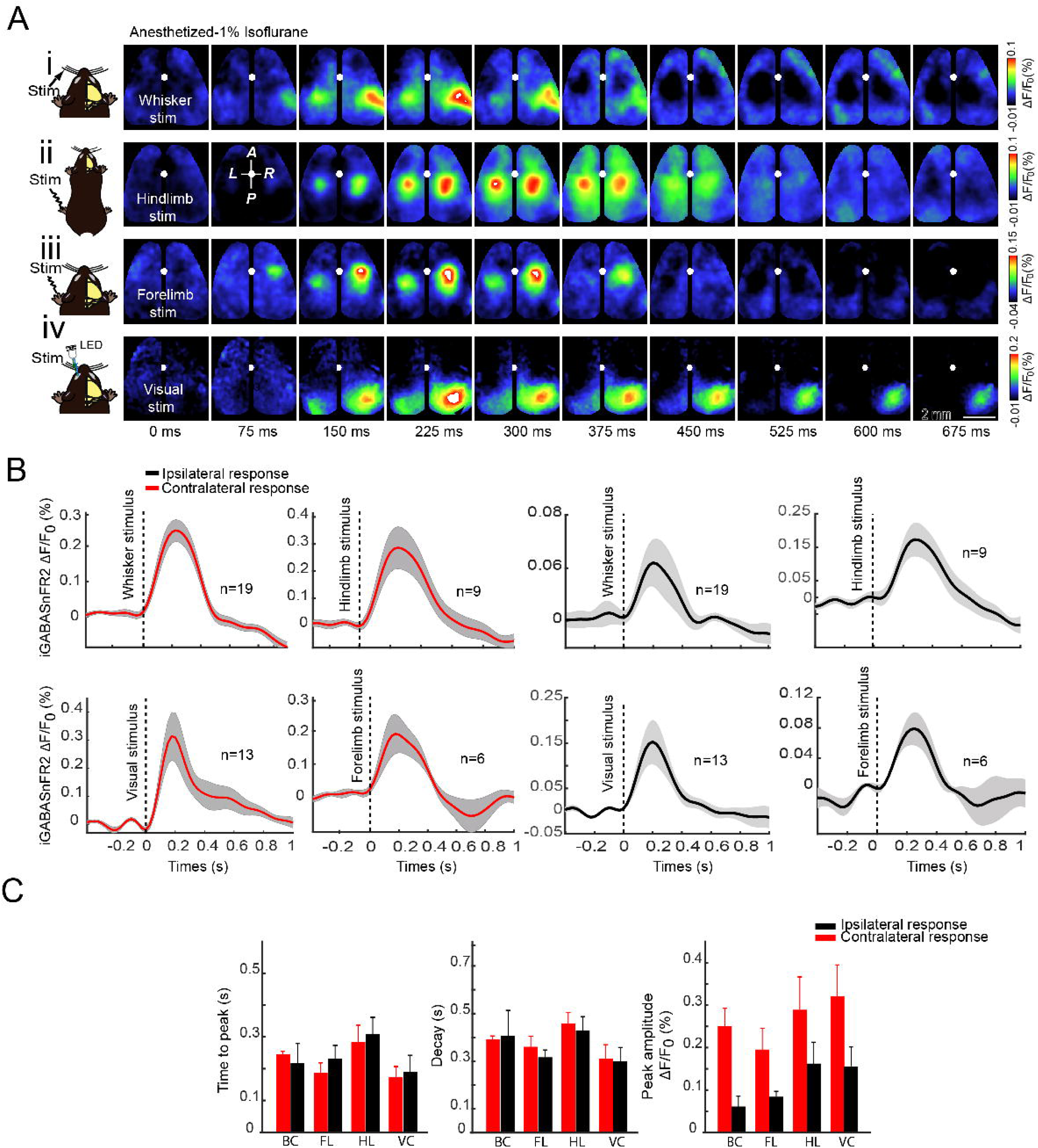
Sensory-evoked GABAergic responses in the neocortex measured by iGABASnFR2 imaging. A. Montages of the wide bilateral craniotomy, with bregma marked by a white circle. Cortical GABAergic activation patterns are shown in a mouse anesthetized with isoflurane (1%) following: (i) whisker stimulation (300 ms), (ii) hindlimb stimulation (300 ms), (iii) forelimb stimulation (300 ms), and (iv) visual stimulation (20 ms) of the eye using an LED. Sensory-evoked extracellular GABA signals were detected using the iGABASnFR2 sensor. Activation is observed within 50–375 ms post-primary sensory cortex activation. Responses represent an average of 40 trials. The second image in the second row indicates anterior (A), posterior (P), medial (M), and lateral (L) directions. B. Time series of sensory-evoked GABA responses. The time series of GABA responses for each sensory stimulation was measured from the respective primary sensory regions. Contralateral responses are shown in black, and ipsilateral responses are shown in red. Data are presented as mean ± SEM, with responses extracted from 3 × 3-pixel regions of interest (ROI) (∼40,401 µm²), n = number of animals. C. Summary of sensory-evoked GABA response features. Decay time (ms), Peak amplitude (ΔF/F) and Time to peak (ms) for contralateral and ipsilateral responses. Data are shown as mean ± SEM, with contralateral responses in blue and ipsilateral responses in red. Statistical Analysis (Two-Way ANOVA): A two-way ANOVA revealed a significant effect of Region on decay time (p = 0.0479), peak amplitude (p = 0.0048), and time to peak (p < 0.0001), indicating sensory-region-dependent differences in GABA responses. Laterality (contralateral vs. ipsilateral) significantly affected peak amplitude (p = 0.0076) but not decay time (p = 0.9874) or time to peak (p = 0.5535). A significant Region × Laterality interaction was found for time to peak (p = 0.0055), indicating lateralization effects vary across sensory regions.

### Sensory-evoked and spontaneous GABA activity in quiet wakefulness resembles anesthesia-induced patterns

Cortical brain states vary across behavioral conditions, shaping spontaneous activity and sensory processing. During active behavior, such as locomotion or whisking, cortical activity becomes desynchronized, and inhibition is modulated to refine sensory gain^47–49^. In contrast, during quiet wakefulness and under light anesthesia, neuronal activity is dominated by slow, synchronized fluctuations that reflect reduced arousal and a shift toward global inhibitory tone^43,50^. While previous studies have shown that GABAergic interneurons contribute significantly to these state-dependent dynamics^36,51^, it remains unclear whether the spatiotemporal profile of extracellular GABA during quiet wakefulness resembles that observed under anesthesia. To examine whether GABAergic responses to sensory stimulation and spontaneous activity in quiet awake mice exhibit spatiotemporal dynamics similar to those observed under anesthesia, we performed wide-field imaging of the cortex using the iGABASnFR2 sensor. Head-fixed mice were imaged in both quite awake and anesthetized states, the latter induced by 1% isoflurane. EMG recordings from a neck muscle electrode, along with video monitoring of body and whisker movement, were used to classify behavioral state (**Figure 3A).** Power spectral analysis of EMG signals confirmed a reduction in muscle tone under anesthesia compared to the quiet awake state. We recorded cortical GABA signals evoked by contralateral visual or whisker stimulation, averaging responses across 40 trials for each condition (**Figure 3B**). Both anesthetized and quiet awake states showed robust stimulus-evoked increases in extracellular GABA in primary sensory areas, including the primary visual cortex (VISp), anterior visual area (VISa), barrel cortex (BC), and primary motor cortex (M1). In Figure 3Bi, which shows cortical GABA responses to visual stimulation, the quiet awake state is characterized by an earlier onset and more widespread GABA release across the visual cortex. In contrast, under anesthesia, the GABA signal appears later and is more spatially restricted. Similarly, in Figure 3Bii, following whisker stimulation, the quiet awake state shows strong and broad activation of the contralateral barrel cortex (BC) and associated motor areas (M1). This response is both faster and more spatially extensive than in the anesthetized state. To assess how GABAergic responses vary across brain states, we analyzed their temporal dynamics during visual and whisker stimulation (**Figure 3C**). Responses were stronger and more spatially distinct during quiet wakefulness than under anesthesia—VISp > VISa for visual input and BC > M1 for whisker input. Quantitative analysis (Figure 3D) confirmed significantly higher peak amplitudes and longer decay times in the awake state.Time to peak was generally shorter in the awake state, suggesting faster inhibitory onset, though most differences were not statistically significant. However, BC in wakefulness responded significantly faster than M1 under anesthesia. Decay times were longer under anesthesia, especially in BC, suggesting more prolonged inhibition when cortical activity is suppressed.

**Figure 3.**
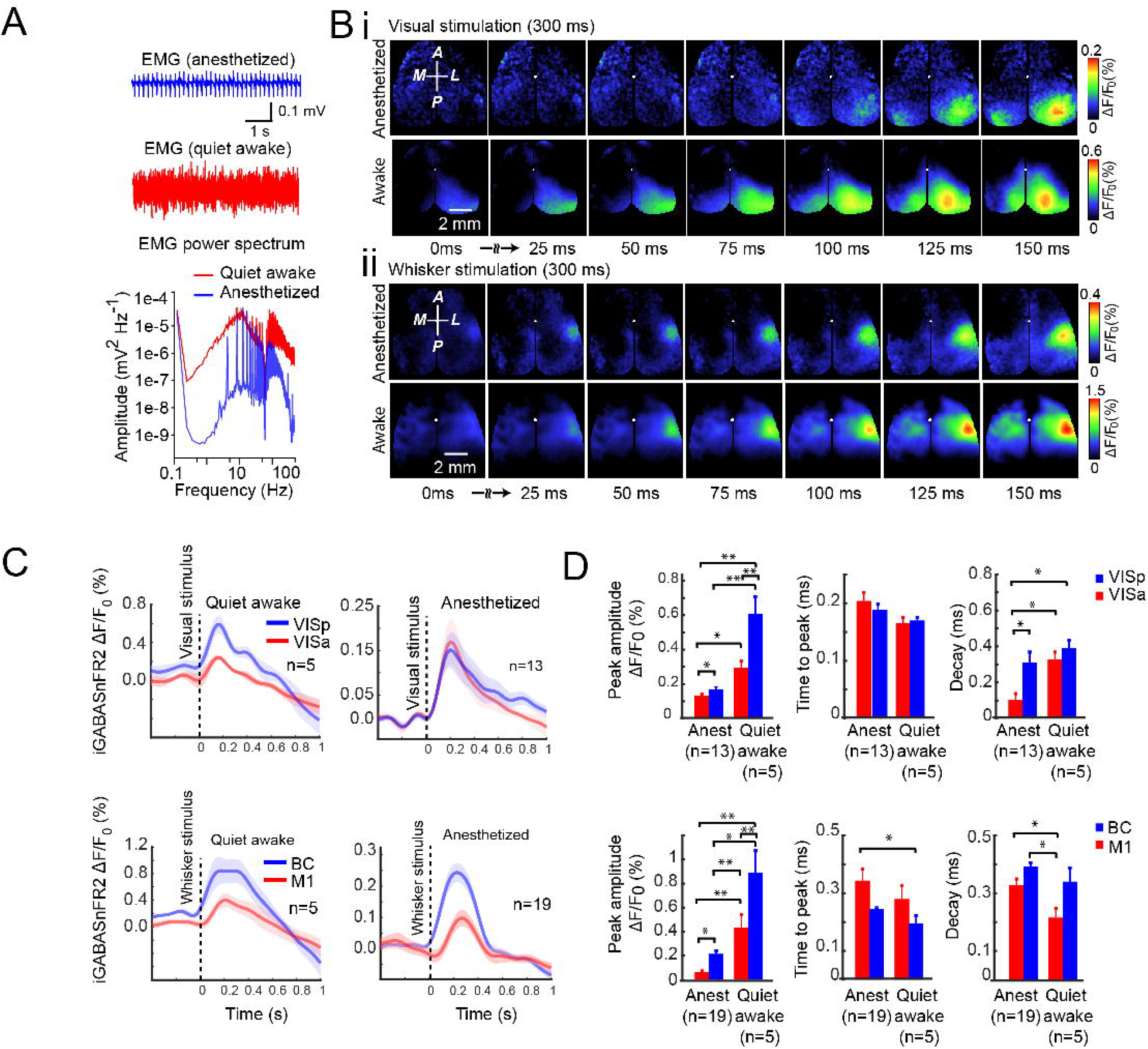
Sensory-evoked and spontaneous GABA activity in quiet wakefulness resembles patterns observed under anesthesia. (A) Individual example of neck muscle EMG (>1 Hz) from a head-restrained mouse under isoflurane (1%) anesthesia (top trace) and quite awake (bottom trace). The power spectra of the EMG signals differ between states of anesthesia (blue) and wakefulness (red). (B) Representative cortical GABA signals are taken from the iGABASnFR2 sensor in response to contralateral visual or whisker stimulation during anesthesia or in quiet awake states. The images represent an average of 40 trials of stimulation (C) Quantification of GABA signals in response to sensory stimulation under anesthesia and quiet wakefulness. Plots show averaged responses from 3 × 3-pixel ROIs (∼0.04 mm²) within VISp (blue) and VISa (red) for visual stimulation and BC (blue) and M1 (red) for whisker stimulation (Ciii, Civ). Shaded areas represent SEM. (D) Statistical comparison of peak amplitude, time to peak, and decay time of GABA responses across states. *P < 0.05, **P < 0.01, one-way ANOVA. Error bars indicate SEM.

These results highlight the influence of both brain state and region on the strength and timing of GABAergic responses.

### Mesoscale imaging of cortical GABA dynamics during natural sleep and wakefulness

While many studies have explored cortical dynamics across sleep and wake states using electrophysiological methods and excitatory activity sensors^52–54^, the ability to track inhibitory signaling at mesoscale resolution across natural brain states remains limited. To further assess iGABASnFR2 performance across different brain states, we examined cortical GABA dynamics during quiet wakefulness, non-rapid eye movement (NREM) sleep, and rapid eye movement (REM) sleep. Mesoscale iGABASnFR2 imaging was combined with simultaneous hippocampal LFP and EMG recordings in head-fixed mice. Animals were allowed to transition naturally between vigilance states while cortical GABA levels were monitored (Figure 4A–C). GABA signals were highest during wakefulness and reduced during NREM sleep (Figure 4D). Spectral analysis showed a reduction in low-frequency GABA fluctuations during REM compared to both wakefulness and NREM (Figure 4E), suggesting diminished slow GABA oscillations during REM sleep. To further assess spatial coordination of cortical GABA activity, we computed pairwise correlation maps across the cortex during each brain state over multiple days. As shown in (**Figure 4F**), Cortical GABA signals exhibited strong bilateral synchrony during quiet wakefulness, which was markedly reduced during NREM sleep and only moderately diminished during REM. This pattern is evident in interhemispheric correlation heatmaps (Figure 4F), where NREM shows the most substantial decrease in bilateral synchrony, while REM correlations remain relatively higher. Quantitative analysis confirmed that mean interhemispheric correlation values were lower during NREM sleep compared to both wakefulness and REM (Figure 4G). During transitions from REM to wakefulness, we observed a sharp increase in cortical GABA levels (Figure 4H). This transition was accompanied by a broad re-engagement of cortical GABAergic activity across multiple regions (**Figure 4I**), suggesting rapid reinstatement of GABA response upon arousal from REM sleep. A detailed overview of the motion-based and electrophysiological features used for behavioral state classification is provided in **Supplementary** Figure 5, which illustrates the temporal alignment of whisker pad and nose motion, EMG power, body motion, theta-to-delta ratio, and hippocampal LFP signals across NREM and REM sleep transitions. Together, these findings demonstrate that iGABASnFR2 reliably captures spontaneous, brain state-dependent fluctuations in extracellular GABA with high temporal and spatial resolution. It supports its utility for long-term mesoscale imaging of cortical inhibition under natural physiological conditions.

**Figure 4.**
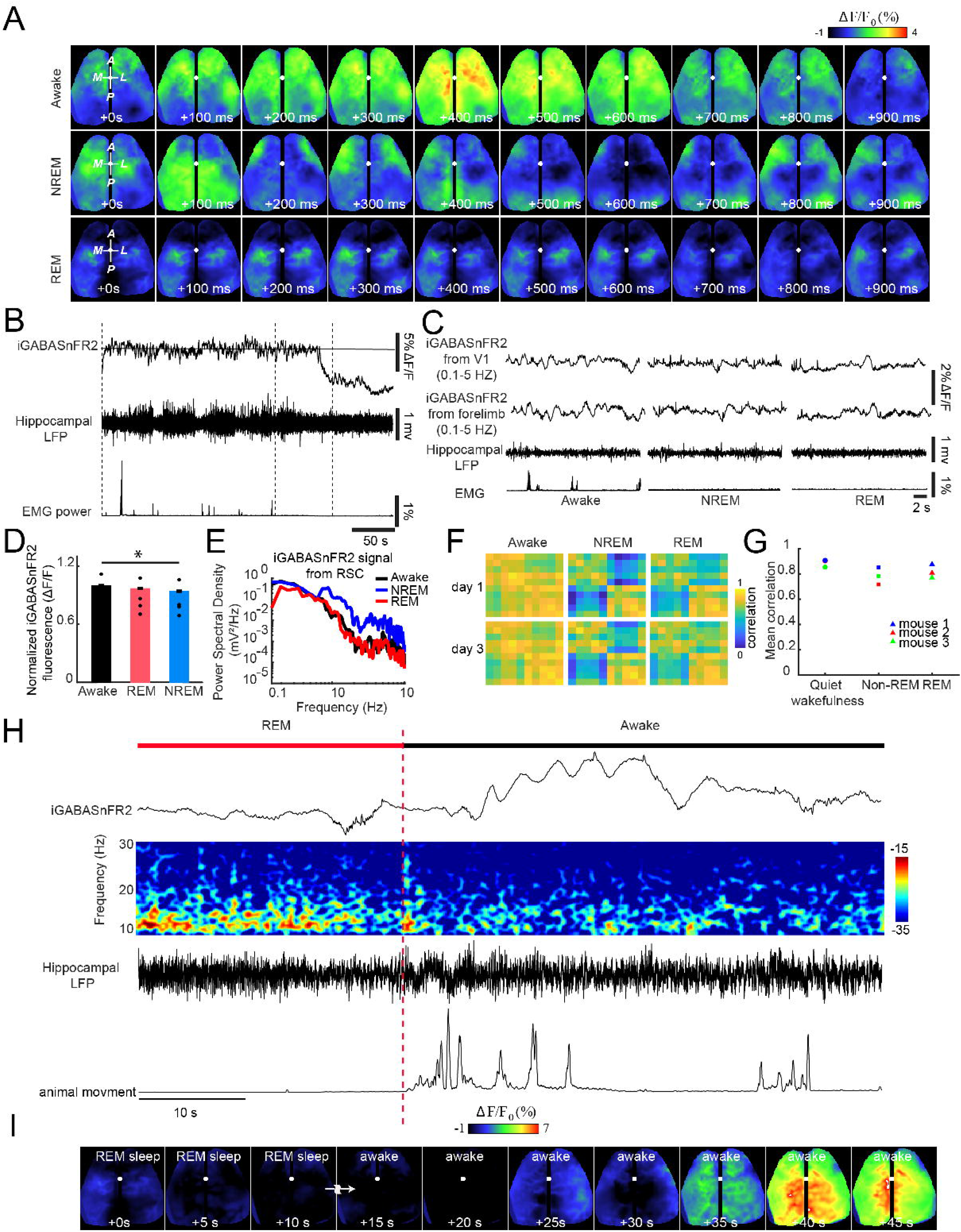
Combined electrophysiological recording and mesoscale iGABASnFR2 imaging of GABA activity during wakefulness and sleep. Spatiotemporal dynamics of GABA activity over a 1-s period during wakefulness, NREM, and REM sleep. Scale bar, 2 mm. (B) Representative traces of GABA activity (RSC), hippocampal LFP, and EMG power during wakefulness, NREM, and REM sleep in a head-fixed mouse. Baseline fluorescence (F) was calculated as the mean signal over the recording session. (C) Expanded view of GABA activity, LFP, and EMG power corresponding to the time windows in (B). (D) Group mean normalized GABA signal across wakefulness, NREM, and REM sleep (n = 5 mice, Kruskal-Wallis test with Nemenyi post hoc correction; *P* = 0.003 overall, all pairwise comparisons significant at *P* < 0.05) (E) Spectral power of GABA signal in retrosplenial cortex (RSC) across wakefulness, NREM, and REM sleep. (F) Cortical GABA activity correlation maps across quiet wakefulness, NREM, and REM sleep over two recording days. **(G)** Mean interhemispheric correlation of cortical GABA activity across wakefulness, NREM, and REM sleep (n = 3 mice). Mean correlation between units significantly increased from quiet wakefulness to NREM and REM sleep across all mice (*p* < 0.05, repeated measures ANOVA with Bonferroni-corrected paired t-tests). **(H)** Simultaneous recordings of cortical GABA signals, hippocampal LFP spectrogram, and EMG power during a REM-to-wakefulness transition. **(I)** Time-lapse montage showing cortical GABA dynamics during the REM-to-wake transition shown in (H). Scale bar: 2 mm.

### Intracortical long-range GABAergic correlations revealed by seed-pixel analysis across brain states

To assess the organization of spontaneous extracellular GABA dynamics in the cortex, we used seed-pixel correlation mapping of iGABASnFR2 fluorescence to investigate intracortical long-range connectivity across different brain states. By placing seed pixels in primary sensory regions, we generated correlation maps of spontaneous GABA dynamics across awake, NREM, and REM sleep states (Figure 5A). In the awake state, these maps revealed strong bilateral synchrony and widespread long-range connectivity between distant cortical regions. These spatial patterns of functional connectivity closely resemble those previously observed with excitatory signals using iGluSnFR and Ca² imaging, suggesting that spontaneous GABA activity also reflects underlying anatomical connectivity and shared network drive^24,25,30^. During NREM sleep, seed-pixel correlations were reduced, indicating a decoupling of large-scale inhibitory networks consistent with cortical slow-wave activity. In contrast, REM sleep preserved many of the bilateral and local connections seen in wakefulness, although with moderate reductions in correlation strength. We further quantified interhemispheric connectivity using pairwise correlation matrices of bilateral cortical regions (**Figure 5B**). These patterns, consistently captured using iGABASnFR2, highlight the sensor’s sensitivity to state-dependent fluctuations in extracellular GABA and its utility for mapping mesoscale inhibitory networks in vivo.

**Figure 5.**
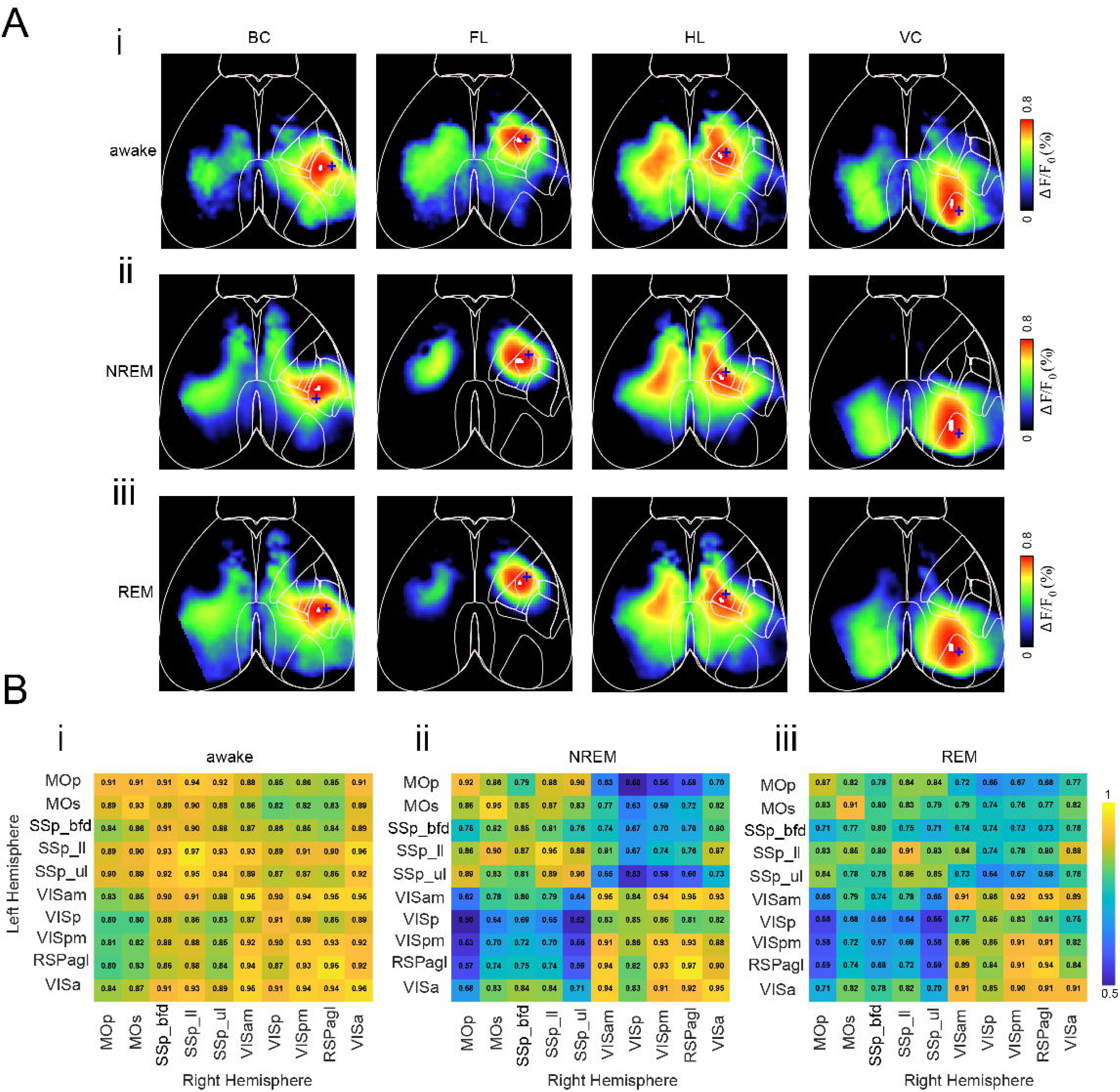
Brain state–dependent patterns of extracellular GABA dynamics measured by iGABASnFR2. (A) Seed-pixel correlation maps (0.1–5Hz) of spontaneous iGABASnFR2 fluorescence across brain states. (i) Awake, (ii) NREM sleep, and (iii) REM sleep. Correlation maps are shown for seed regions in the barrel cortex (BC), forelimb cortex (FL), hindlimb cortex (HL), and visual cortex (VC). (B) Interhemispheric correlation matrices of spontaneous extracellular GABA signals across brain states. (i) Awake, (ii) NREM sleep, and (iii) REM sleep. Matrices display Pearson correlation coefficients between bilateral cortical regions based on iGABASnFR2 fluorescence. Regions include MOp – Primary motor area, MOs – Secondary motor area, SSp_bfd – Primary somatosensory area (barrel field), SSp_ll – Lower limb, SSp_ul – Upper limb, VISam – Anteromedial visual area, VISp – Primary visual area, VISpm – Posteromedial visual area, RSPagl – Agranular retrosplenial area, VISa – Anterior visual area. Widespread interhemispheric connectivity is observed during wakefulness and REM sleep, with reduced correlations during NREM sleep. Blue crosses indicate the location of the seed pixel used for correlation mapping. number of animals=5.

### Tiagabine elevates baseline GABA levels but dampens sensory-evoked responses and reorganizes cortical inhibitory connectivity

To examine how pharmacological inhibition of GABA reuptake influences cortical GABA dynamics, we administered Tiagabine—a selective GAT-1 inhibitor^55^—under 1% isoflurane anesthesia and monitored extracellular GABA levels using iGABASnFR2. Mice were head-fixed throughout the experiment. We first recorded visually evoked GABA responses (∼6 minutes), followed by a ∼15-minute baseline period of spontaneous activity. While the mouse remained head-fixed, Tiagabine was injected intraperitoneally. Spontaneous activity was then recorded for another ∼15 minutes before delivering a second round of visual stimulation to assess post-Tiagabine responses. Before Tiagabine injection, visual stimulation evoked robust and spatially localized increases in GABA signals within the contralateral primary visual cortex (Figure 6Ai). Following Tiagabine administration, the same sensory stimulus failed to evoke any detectable response (Figure 6Aii). This complete loss of evoked GABA activity is also evident in the time-series traces (Figure 6Aiii). These recordings were acquired at 150 Hz using the blue fluorescence channel under continuous (non-strobing) illumination.

**Figure 6.**
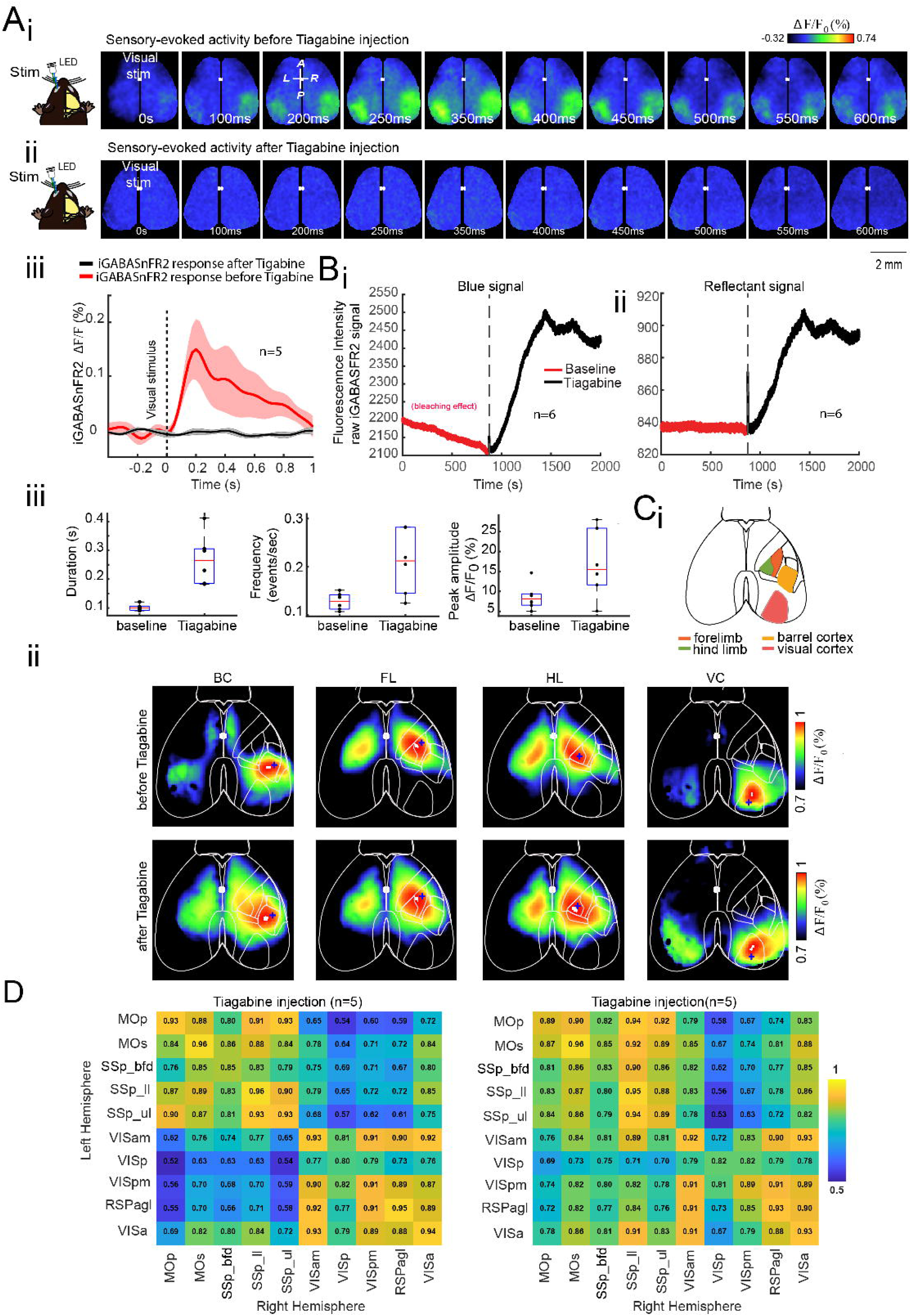
Effect of Tiagabine administration on extracellular GABA activity in mice expressing iGABASnFR2. **(A)** Sensory-evoked GABA responses recorded with the blue channel (iGABASnFR2) at 150 Hz under continuous illumination (no strobing). **(i)** Representative montages of visually evoked GABAergic activity before Tiagabine administration. **(ii)** The same stimulation after Tiagabine injection. **(iii)** Time series of contralateral visually evoked iGABASnFR2 signals from the primary visual cortex (VISp). Traces represent mean ± SEM from 5 animals, extracted from 3 × 3-pixel ROIs (∼0.04 mm²). **(B)** Tiagabine-induced changes in spontaneous GABA activity in VISp, recorded using dual-channel strobing acquisition (80 Hz total: 40 Hz blue, 40 Hz green). **(i)** Blue fluorescence signal (iGABASnFR2) from VISp, averaged across 6 animals. Traces show signal before (red) and after (black) Tiagabine injection. The red trace shows a photobleaching trend; the black trace shows increased fluorescence post-injection. **(ii)** Reflectance signal from the same VISp region, averaged across 6 animals. **(iii)** Quantification of spontaneous iGABASnFR2 transients in VISp before and after Tiagabine injection. Box plots show duration (left), frequency (middle), and peak amplitude (right) of fluorescence transients. Data represents mean ± SEM across 6 animals (n = 6). Tiagabine significantly increased the duration, frequency, and amplitude of spontaneous GABAergic transients compared to baseline. Statistical comparisons were made using paired t-tests; all changes were significant (*p* < 0.05). **(iv)** Schematic of cortical regions of interest (ROIs) used for seed-pixel correlation analysis: barrel cortex (orange), forelimb (red), hind limb (green), and visual cortex (brown). **(C)** Functional connectivity of spontaneous iGABASnFR2 activity, recorded with dual-channel strobing (40 Hz blue, 40 Hz green). **(i)** Seed-pixel correlation maps (0.1–5 Hz) before and after Tiagabine injection. Blue asterisks indicate seed pixel locations in the selected ROIs. **(ii)** Interhemispheric correlation matrices of spontaneous iGABASnFR2 activity before and after Tiagabine injection (*n* = 5 mice), showing enhanced bilateral cortical connectivity post-injection. Regions: MOp – Primary motor area, MOs – Secondary motor area, SSp_bfd – Primary somatosensory area (barrel field), SSp_ll – Lower limb, SSp_ul – Upper limb, VISam – Anteromedial visual area, VISp – Primary visual area, VISpm – Posteromedial visual area, RSPagl – Agranular retrosplenial area, VISa – Anterior visual area. The interior is at the top of all maps. Blue crosses indicate the location of the seed pixel used for correlation mapping.

To better understand this loss of evoked responsiveness, we analyzed spontaneous GABA activity using dual-wavelength strobing imaging (40 Hz blue, 40 Hz green). In the primary visual cortex, the raw blue fluorescence signal, averaged across 6 animals, showed a sustained elevation^56–57^ in baseline GABA levels following Tiagabine injection (Figure 6Bi), consistent with extracellular GABA accumulation due to GAT-1 blockade. Reflectance signal from the same region also increased post-injection (Figure 6Bii), confirming overall signal elevation.

Quantitative analysis revealed that Tiagabine significantly increased the duration, frequency, and peak amplitude of spontaneous GABA transients across animals (Figure 6Biii), reflecting heightened inhibitory tone.

To assess how elevated GABA levels affect network organization, we performed seed-pixel correlation analysis (0.1–5 Hz) of spontaneous iGABASnFR2 activity. Following Tiagabine injection, seed regions in sensory cortices showed increased local synchrony and expanded spatial correlation patterns (Figure 6C). Interhemispheric correlation matrices also revealed stronger bilateral connectivity, particularly between homologous sensory areas (Figure 6D). These results further validate the ability of iGABASnFR2 to detect changes in extracellular GABA levels under both baseline and stimulated conditions, highlighting its sensitivity to dynamic alterations in cortical inhibition and its potential utility in pharmacological studies targeting GABAergic signaling.

## Discussion

We used mesoscale imaging with the genetically encoded GABA sensor iGABASnFR2 to map the spatiotemporal dynamics of extracellular GABA across the intact mouse cortex in vivo. By characterizing extracellular GABA dynamics in both sensory-evoked and spontaneous across brain states—including anesthesia, quiet wakefulness, NREM, and REM sleep—we provide, for the first time, a comprehensive mesoscale imaging of GABA dynamics in the cortex. our results establish iGABASnFR2 as a valuable tool for investigating GABAergic tone and state-dependent fluctuations in extracellular GABA, complementing previous studies that used iGluSnFR and voltage-sensitive dyes to map excitatory signaling and depolarization across large-scale cortical networks^19,23^. The robust and state-dependent changes in extracellular GABA we observed— during sensory stimulation, spontaneous activity, and pharmacological manipulation— demonstrate that inhibitory tone is dynamically regulated across brain states and cortical regions.

### Interpreting iGABASnFR2 signals

iGABASnFR2 fluorescence changes are likely shaped by a combination of GABA release from interneurons, diffusion through the extracellular space, and clearance by GABA transporters, particularly GAT-1^58–60^. The timing of sensory-evoked responses—characterized by a delayed onset (∼75 ms) and prolonged signal lasting several hundred milliseconds—is consistent with the dynamics of SST-positive interneuron-mediated inhibition, which is known to follow the rapid activation of PV+ interneurons^41,43^. The prolonged decay phase of iGABASnFR2 signals (**Figure 2C**) suggests that extracellular GABA may persist longer, potentially contributing to modulatory or volume transmission effects^61^.

### Cortical GABA responses to sensory input are conserved across brain states

Sensory-evoked results revealed robust, modality-specific patterns of extracellular GABA activation across the cortex, reflecting both localized responses and broader, network-level dynamics (**Figure 2A**). Each stimulus modality selectively activated its corresponding primary sensory area (**Figure 2B**), with activity spreading from the focal point of stimulation to more distal, functionally connected cortical regions. This widespread response pattern suggests that inhibitory activity can propagate across large-scale networks, challenging the traditional view of inhibition as strictly local.

In our study, contralateral responses were consistently stronger and faster than ipsilateral responses, likely reflecting direct thalamocortical projections to the primary sensory cortex. In contrast, the delayed and weaker ipsilateral responses are consistent with slower callosal transmission and interhemispheric integration (Figure 2B–C), as reported in previous studies of cortical sensory processing^45^. In addition, despite differences in global cortical state and arousal, sensory-evoked GABAergic responses were consistently observed across brain states, suggesting a preserved functional role of inhibition, even as response strength and timing are modulated (Figure 3B–D). This robustness underscores the reliability of iGABASnFR2 for monitoring inhibitory dynamics and reflects the essential role of inhibition in stabilizing excitation and regulating cortical gain, as also supported by prior calcium imaging studies in awake animals ^47^.

### Spontaneous GABA dynamics reveal state-dependent connectivity

In our study, interhemispheric GABA synchrony was lowest during NREM sleep (Figure 4D-G), suggesting that inhibition in this state becomes locally structured and functionally decoupled across hemispheres. REM sleep (Figure 4D-G) showed partial recovery of interhemispheric GABA synchrony, suggesting a reorganization of inhibitory networks distinct from both wakefulness and NREM. This contrasts with vascular-based findings; for instance, a study ^62^reported strong bilateral hemodynamic coherence during both NREM and REM, reflecting broader, nonspecific signals. However, they also noted that neurovascular signals integrate contributions from neurons, astrocytes, and metabolism, potentially decoupling them from electrophysiological activity. Our direct measurement of extracellular GABA provides a distinct view, revealing that strong vascular synchrony during NREM does not imply coordinated inhibitory signaling. Indeed, the low GABA synchrony we observe during NREM contrasts with the high bilateral [HbT] coherence reported by Turner and colleagues ^62^, highlighting a dissociation between vascular and inhibitory network dynamics.

Similar patterns have been observed in BOLD fMRI studies, which report reduced large-scale connectivity during NREM and partial restoration during REM^63–65^. These converging lines of evidence highlight arousal state as a key factor in shaping both vascular and neural network dynamics and emphasize the central, yet state-specific, role of inhibition in large-scale brain coordination during sleep.

### Functional inhibitory architecture aligns with cortical structural organization

Seed-pixel correlation results revealed structured long-range inhibitory connectivity (**Figure 5A**). In the awake state, we observed strong bilateral synchrony between homologous sensory areas and strong local correlations within modalities (**Figure 5Ai, 5Bi**). These maps are similar to those derived from excitatory indicators, including iGluSnFR and VSD^19,23,66^, supporting the view that inhibitory and excitatory networks are functionally integrated, but exhibit distinct temporal dynamics^42^. In NREM sleep, long-range inhibitory connectivity weakened, suggesting a shift toward localized processing (**Figures 5Aii, 5Bii)** while REM sleep partially restored (**Figures 5Aiii, 5Biii**). State-dependent changes highlight the flexibility of cortical inhibitory networks, which adapt dynamically to behavioral states, promoting broad coordination during wakefulness and preserving localized stability during sleep^67^.

### Tiagabine elevates extracellular GABA and disrupts sensory-evoked responses via altered cortical synchrony

Our data show that Tiagabine disrupts sensory-evoked GABAergic responses by elevating extracellular GABA and altering cortical network dynamics. Before injection, visual stimulation produced robust, spatially confined increases in GABA sensor signals in the visual cortex (Figure 6A). After Tiagabine administration, however, the same stimulus no longer evoked detectable GABA responses—despite a sustained increase in baseline fluorescence (Figure 6Bi), confirming that the sensor remained active. Although not used for quantification, an increase in green reflectance signal (Figure 6Bii) suggested broader changes in cortical vascular or metabolic state. The absence of evoked activity in the presence of elevated GABA points to a nonlinear relationship between inhibitory tone and sensory responsiveness. This may involve excessive tonic inhibition, desynchronized interneuron networks, or saturation of GABAergic circuits. In our study, Tiagabine-induced increases in extracellular GABA were clearly detected by iGABASnFR2, with fluorescence peaking around 10 minutes post-injection—consistent with prior microdialysis reports^68^. This elevation likely enhanced tonic inhibition via extrasynaptic GABA_A receptors^69^, reducing neuronal excitability and suppressing phasic, stimulus-driven responses. This mechanism helps explain the loss of stimulus-locked GABA transients. Similar effects have been observed in rodent EEG studies, where Tiagabine increases low-frequency synchronization and, at higher doses, induces hypersynchronous activity resembling absence seizures—likely a result of widespread tonic inhibition^70^. Our results align with these findings and demonstrate that iGABASnFR2 effectively detects sustained changes in extracellular GABA. However, the loss of evoked transients following Tiagabine suggests the sensor may reach a functional ceiling under conditions of high ambient GABA, limiting its dynamic range. Though Tiagabine does not directly cause transporter reversal, elevated intracellular GABA and transporter saturation could still favor nonvesicular GABA release^67^, further amplifying tonic signaling. Together, these findings highlight both the strengths and limitations of iGABASnFR2: it is sensitive to pharmacologically induced increases in extracellular GABA but may underestimate fast, transient responses when tonic levels are elevated. This underscores the importance of considering inhibitory tone and network state when interpreting GABA imaging data.

## Conclusion

Our findings establish iGABASnFR2 as a robust sensor for widefield imaging of extracellular GABA in the cortex, revealing both strong stimulus-locked responses and state-dependent dynamics. Our experiments show that inhibition is not merely local or reactive but is organized into large-scale motifs that shift with arousal state and can be pharmacologically tuned. For a long time, efforts to understand how excitation and inhibition interact across the cortex have been limited by the lack of tools that can monitor both processes in real time—especially in awake, behaving animals. That’s now changing. Genetically encoded sensors like iGABASnFR2, along with glutamate and calcium indicators for excitatory activity, offer a new window into the excitation-inhibition (E/I) balance with spatial, temporal, and cell-type specificity. Importantly, the development of red-shifted glutamate sensors, such as R-iGluSnFR ^71^, now makes it possible to image GABA and glutamate simultaneously, opening the door to more complete views of circuit function in vivo.This capability will be essential for understanding how the E/I balance is regulated during normal brain function—and how it becomes disrupted in neurological and psychiatric disorders. Many neurological and psychiatric disorders—such as epilepsy, autism, and schizophrenia—involve disruptions in E/I balance, yet direct measurements of these dynamics in vivo have remained elusive. Simultaneous imaging of glutamate and GABA now enables researchers to observe how excitation and inhibition interact across space and time, how this interaction is modulated by behavioral state, neuromodulators, or genetic risk factors, and how it breaks down in pathological conditions. These insights are critical for developing targeted interventions that restore E/I balance and stabilize network function in disease.

## Acknowledgments

We thank Di Shao and the Animal Welfare Committee at the University of Lethbridge for their support in enabling the animal experiments. We also thank Majid H. Mohajerani for providing access to laboratory resources and equipment during the data collection phase of this project. We thank Nicholas J Michelson and Tim H. Murphy for performing the initial experiment.

## Author contributions

E.R. conceptualized the study. E.R. designed and conducted the main experiments. M.N. contributed to additional experimental procedures. J.K.A. implemented the strobing imaging system. E.R. and S.T. analyzed the data. E.R. prepared all the figures, wrote, and revised the manuscript. All authors have read and agreed to the submitted version of the manuscript.

## Supplementary material

Supplementary material and methods are available.

## The code and data Availability

The code and all data analyzed and used to produce the main findings of this study have been deposited on Dryad. Source data files are available for Figures 2, 3, 5, and 6.

## Biography

**Edris Rezaei** is a Ph.D. candidate in the Canadian Centre for Behavioural Neuroscience at the University of Lethbridge, Canada. He is interested in how inhibition shapes neural activity, particularly its role in cognition and neurological disorders.

**Setare Tohidi** completed her master’s degree at the University of Lethbridge. She is now working as a Data Scientist (Junior Associate) at Synechron in Canada.

**Mojtaba Nazari**□**Ahangarkolaee** completed her master’s degree at the University of Lethbridge, affiliated. He is now working as a Data Scientist at the University of Lethbridge.

**J. Karimi Abadchi** is currently a postdoctoral researcher at the Douglas Research Centre in Montreal. He was a Ph.D. student at the University of Lethbridge.

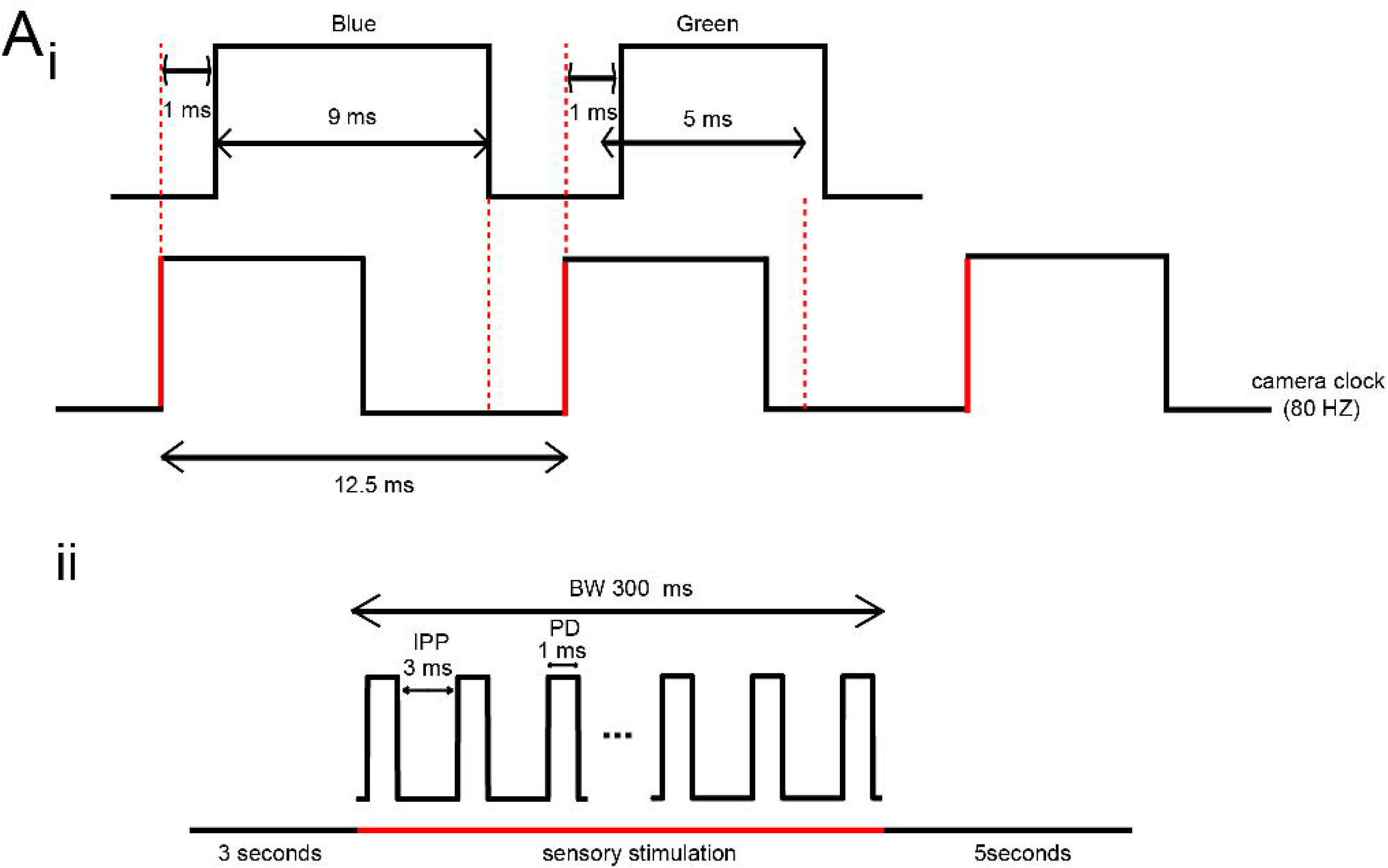

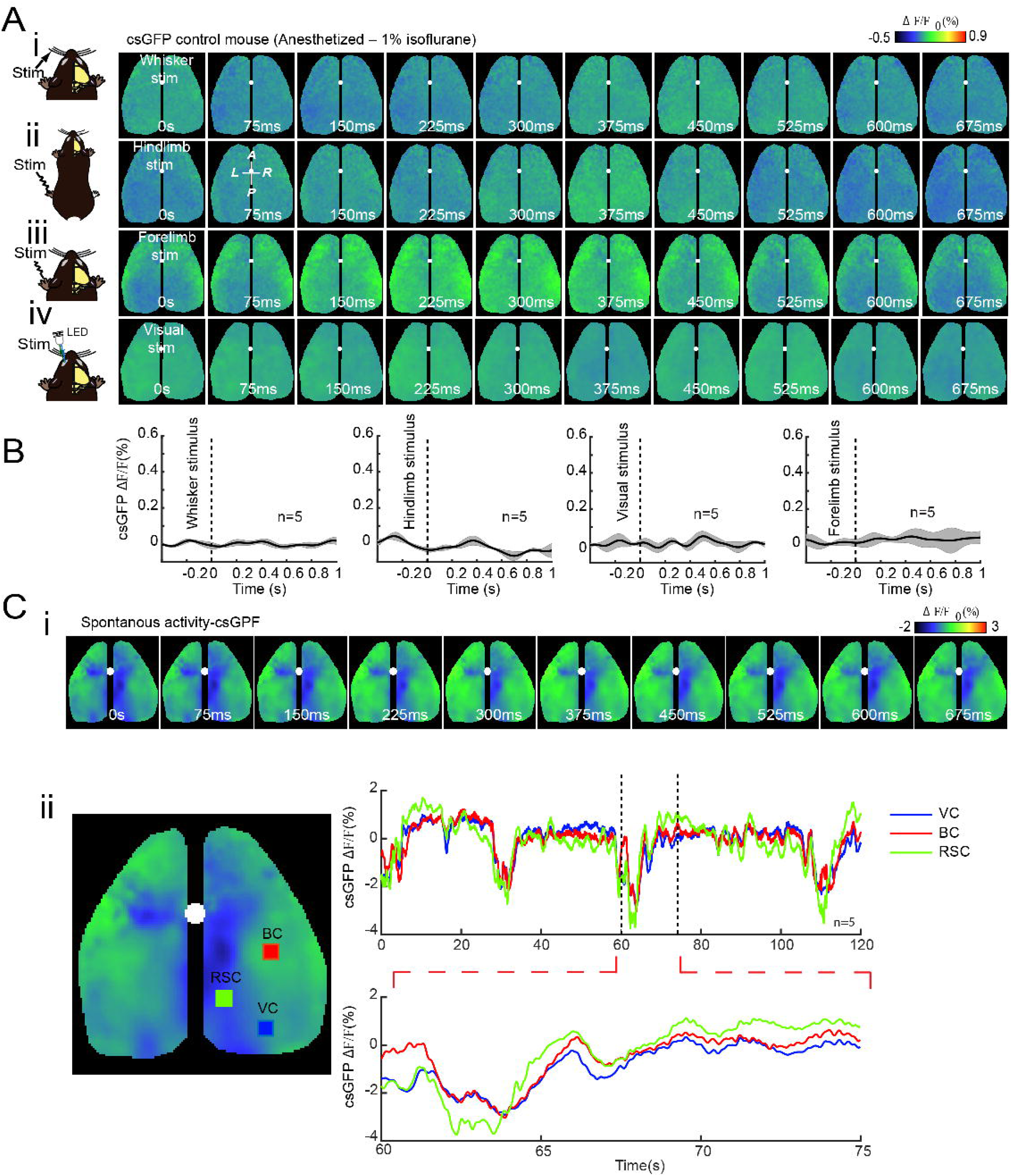

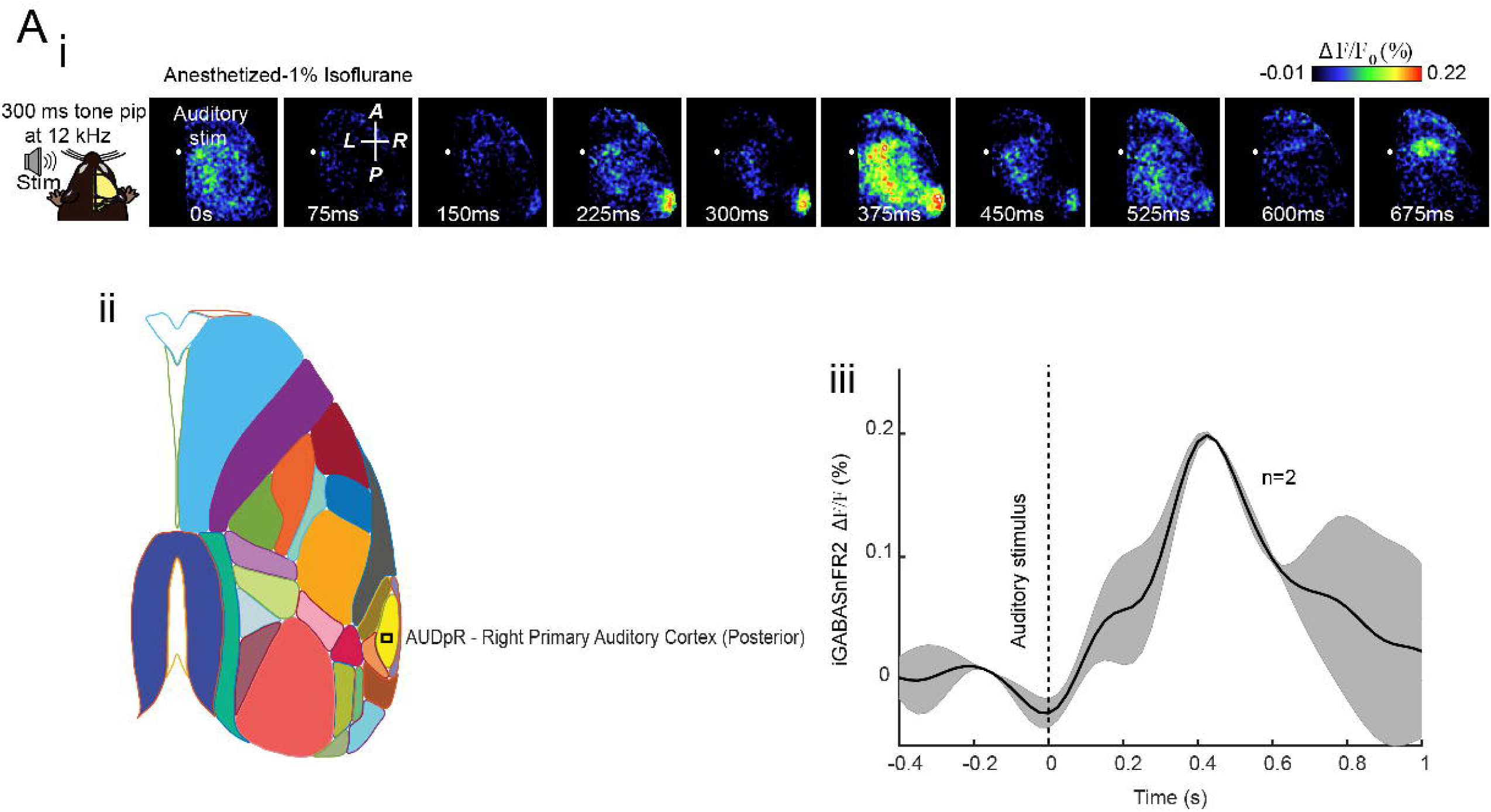

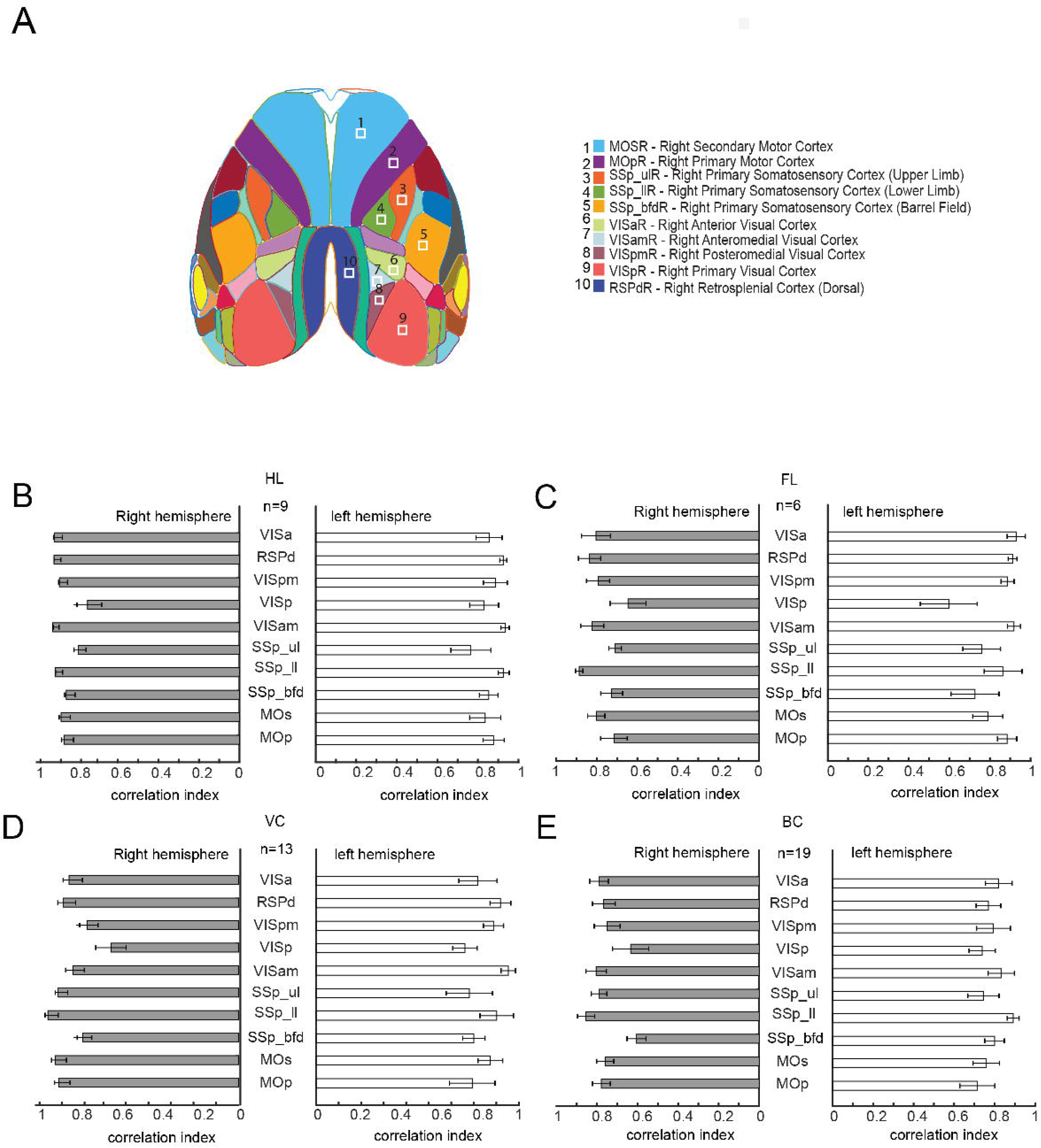

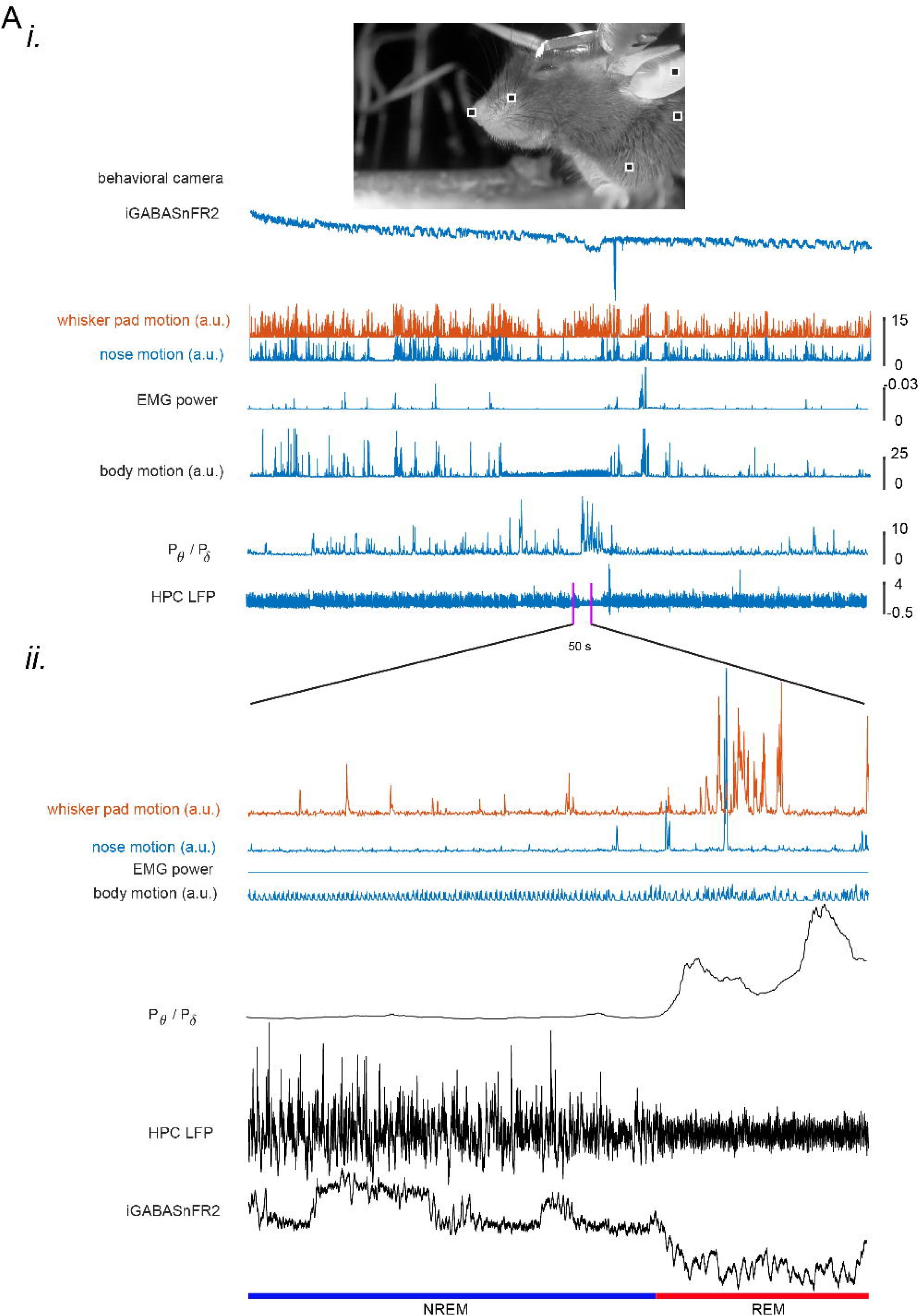

