## Supplementary figures for "Characterization of iGABASnFR2 for in vivo mesoscale imaging of intracortical GABA dynamics"

Edris Rezaei, Setare Tohidi, Mojtaba Nazari-Ahangarkolaee, Javad Karimi Abadchi, Nicholas J Michelson, Timothy H. Murphy

**Supplementary figures 1-6.**

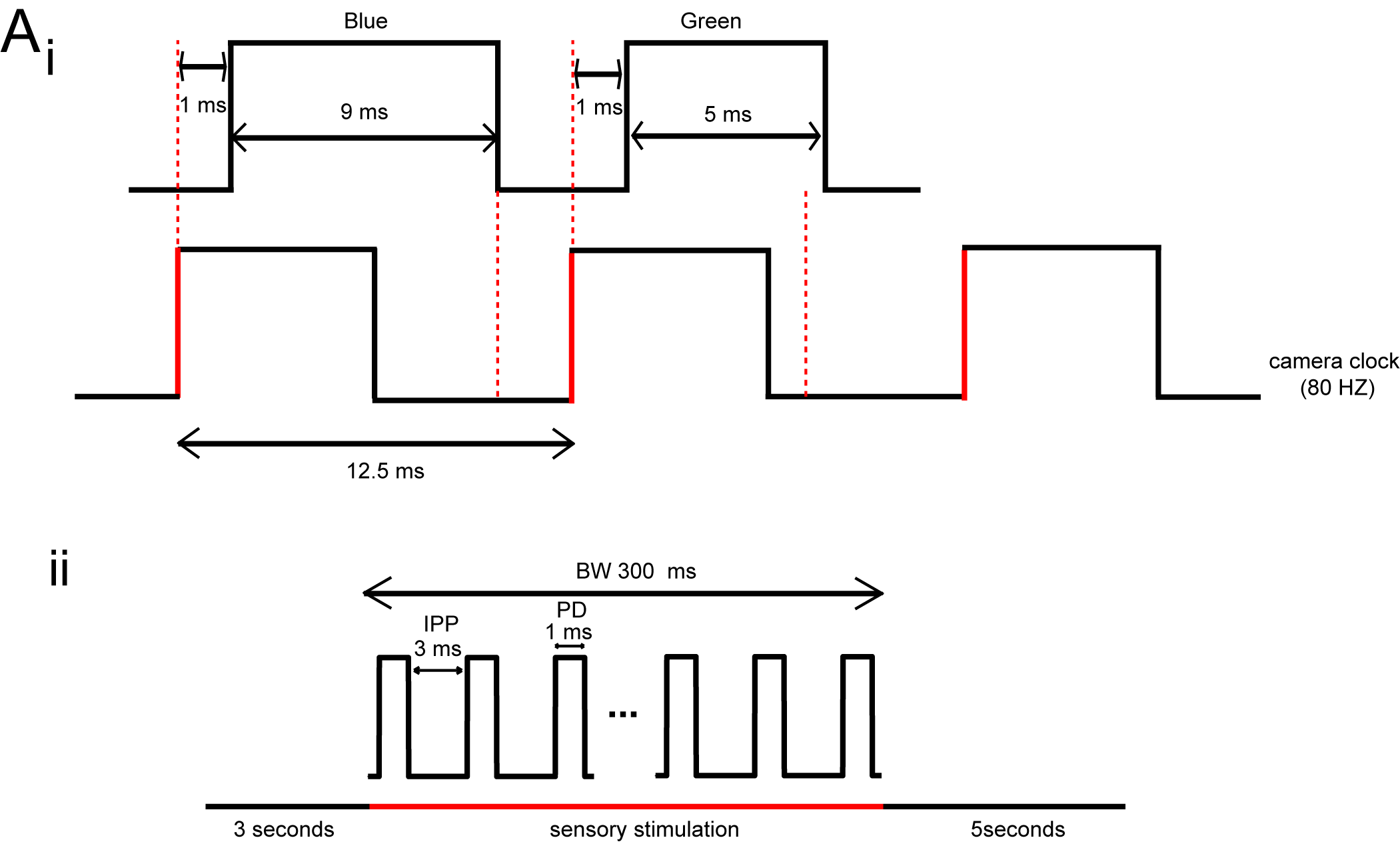
**Supplementary Figure 1. A. Imaging acquisition timing and sensory stimulation protocol.** (i) Diagram illustrating the timing of LED illumination and camera acquisition during dual-channel wide-field imaging at 80 Hz. Top, the timing of blue and green LED activation. Each TTL rising edge (red vertical lines) triggers an imaging frame. After a 1 ms delay, either the blue LED (9 ms ON) or green LED (5 ms ON) is activated, alternating every frame. This design ensures only one LED is active during each ~11 ms camera exposure, preventing spectral contamination. Bottom, the camera clock traces at 80 Hz. Each square pulse represents one frame, with a duration of 12.5 ms, matching the TTL trigger frequency. (ii) Timing diagram of the sensory stimulation protocol. Each trial includes an 8-second acquisition period, with 80 Hz imaging throughout. A piezoelectric actuator delivers mechanical stimulation between 3 and 3.3 seconds in the form of a burst (BW = 300 ms) of pulses (PD = 1 ms) spaced by a 3 ms inter-pulse period (IPP). A total of 40 trials were performed per condition.

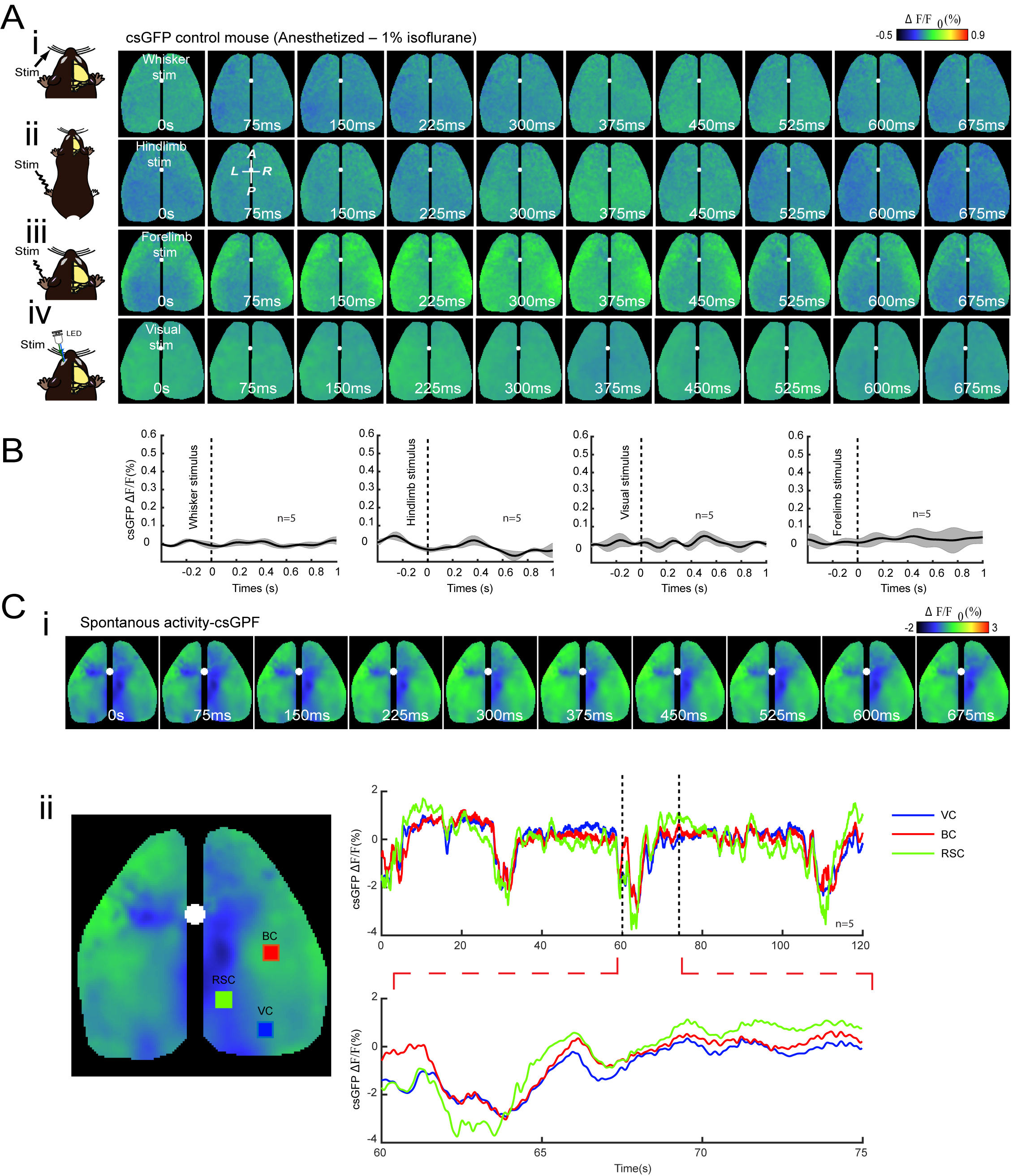

**Supplementary Figure 2. Sensory-evoked and spontaneous cortical responses under anesthesia using csGFP as a control for iGABASnFR2.** (A) Montages showing cortical activation patterns in a mouse anesthetized with 1% isoflurane. A white circle marks Bregma. Sensory-evoked responses are shown for: (i) contralateral whisker (C2) stimulation, (ii) contralateral hindlimb stimulation, (iii) contralateral forelimb stimulation, and (iv) visual stimulation of the contralateral eye using an LED. Sensory responses emerge within 50–375 ms following stimulation, with activation localized to the primary sensory cortex (white arrows). Each response is averaged over 40 trials. The second frame in the second row indicates anterior (A), posterior (P), medial (M), and lateral (L) orientations. (B) Time course plots of ΔF/F (%) for csGFP signals in response to whisker (i), hindlimb (ii), forelimb (iii), and visual (iv) stimulation. Traces represent mean ± SEM from 3 × 3-pixel ROIs (~0.04 mm²); *n* indicates the number of animals. (C) Spontaneous cortical activity measured with csGFP under isoflurane anesthesia. (i) Representative montages showing spontaneous cortical dynamics over time. (ii) Map showing selected ROIs in the right barrel cortex (red), visual cortex (blue), and retrosplenial cortex (orange). Time series plot (top right) displays spontaneous fluorescence fluctuations across 120 seconds. A magnified 15-second window is shown below for detail. Imaging was performed using dual-channel (blue and green) acquisition at 80 Hz, n=number of animals.

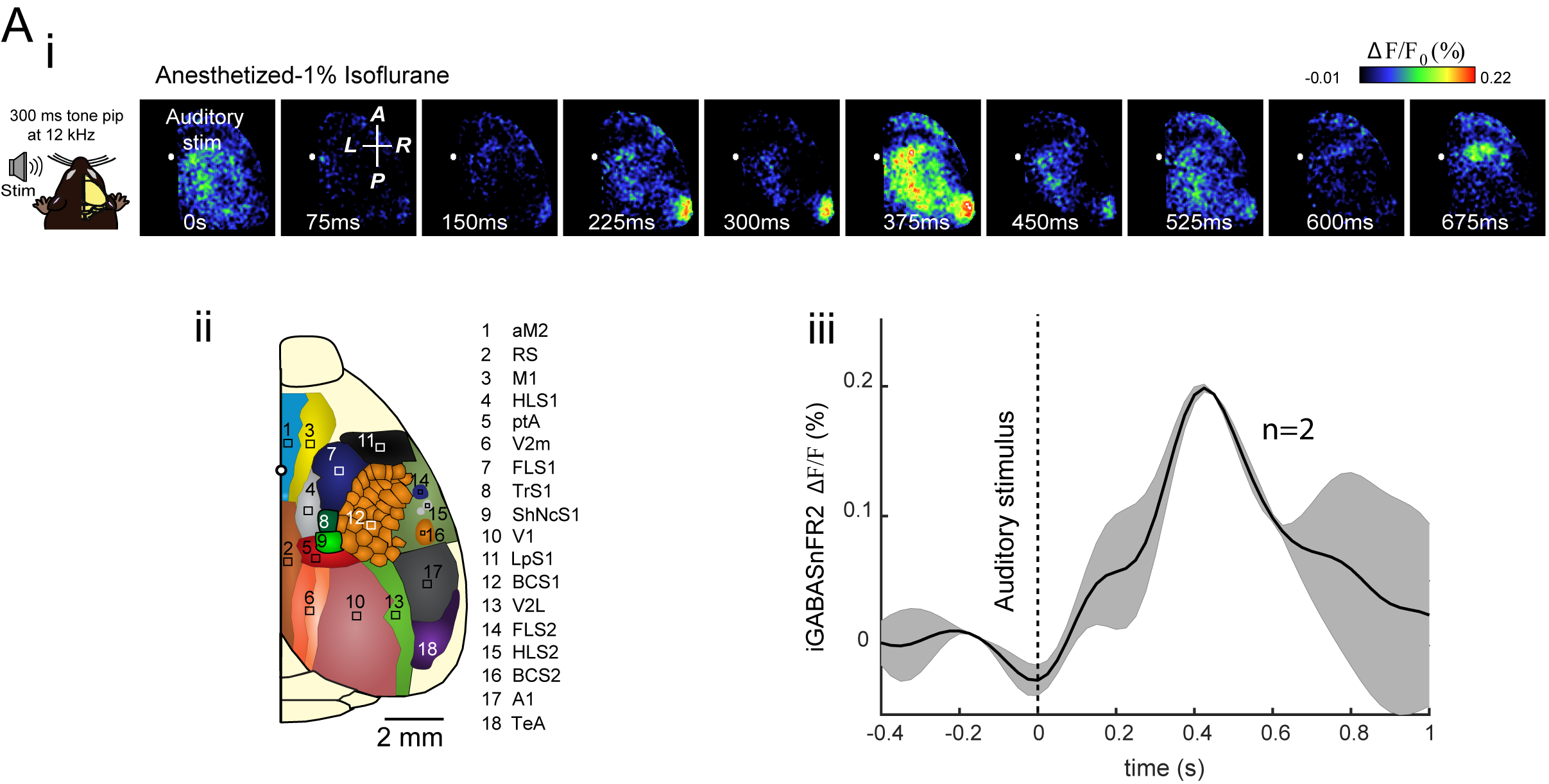
**Supplementary Figure 3. Auditory-evoked cortical GABA responses under anesthesia measured with iGABASnFR2.** (A) GABAergic activity evoked by auditory stimulation in mice anesthetized with 1% isoflurane. (i) Time-lapse montages show changes in iGABASnFR2 fluorescence following 300-ms contralateral auditory stimulation. Images represent the average of 40 trials. The white dot indicates the time of stimulation onset (0 ms). The frame at 75 ms includes a reference compass showing anterior (A), posterior (P), medial (M), and lateral (L) directions. (ii) Allen Mouse Brain Atlas map showing cortical regions used for reference and ROI definition. Regions are labeled with abbreviations and numeric IDs according to atlas convention. The white scale bar represents 2 mm. (iii) Time course of ΔF/F (%) from the contralateral auditory cortex, averaged across two animals (n = 2). Signal traces represent the mean ± SEM extracted from 3 × 3-pixel unilateral regions of interest (~0.04 mm²). The dashed vertical line indicates the onset of auditory stimulation (0 s).

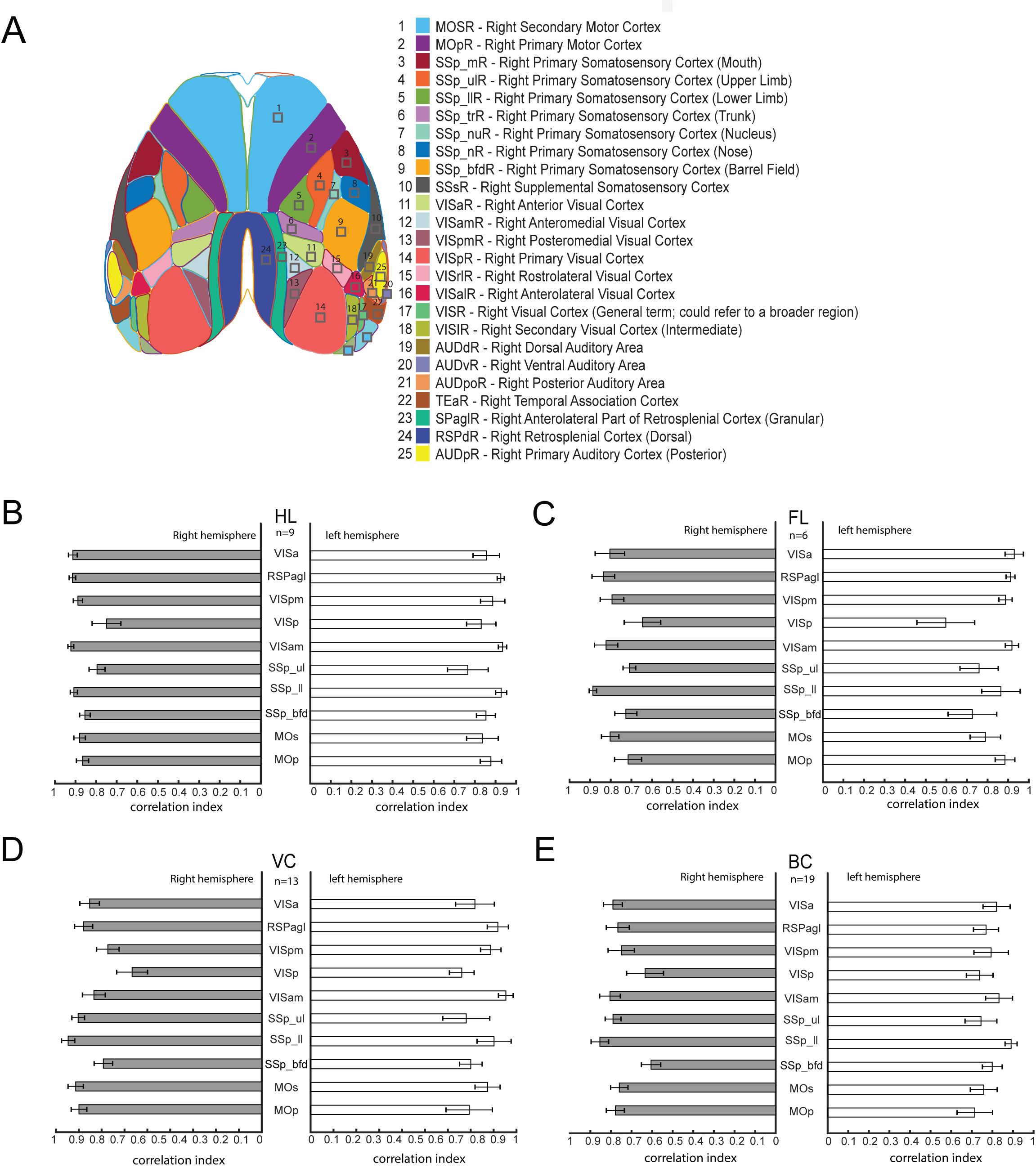

**Supplementary Figure 4. Intrahemispheric connectivity: Region-based comparison of sensory-evoked and spontaneous activity correlation maps.** (A) Schematic of the right hemisphere showing the 10 cortical regions of interest (ROIs) used for intrahemispheric correlation analysis: 1) MOp – Primary motor cortex, 2) MOs – Secondary motor cortex, 3) SSp_bfd – Primary somatosensory area, barrel field, 4) SSp_ll – Primary somatosensory area, lower limb, 5) SSp_ul – Primary somatosensory area, upper limb, 6) VISam – Anteromedial visual area, 7) VISp – Primary visual area (V1), 8) VISpm – Posteromedial visual area, 9) RSPagl – Retrosplenial area, lateral agranular part, 10) VISa – Anterior visual area. To assess intrahemispheric connectivity, we calculated Pearson’s correlation coefficients between the iGABASnFR2 signal from each primary sensory ROI and the 10 target ROIs, based on 3 × 3-pixel regions of interest (~0.04 mm²). These analyses were performed separately for the right and left hemispheres during sensory-evoked conditions. (B–E) Bar plots display the pairwise correlation indices for the hindlimb (HL), forelimb (FL), visual cortex (VC), and barrel cortex (BC), respectively. **Statistical analysis:** Paired two-tailed *t*-tests were used to compare the correlation indices between the right and left hemispheres for each ROI. Bonferroni correction was applied to adjust for multiple comparisons across 10 ROIs. In addition, a two-way repeated-measures analysis of variance (ANOVA) was performed using a linear mixed-effects model with the fixed effects of (the right vs left hemispheres), *ROI*, and the *Condition × ROI* interaction. The degrees of freedom for marginal tests were computed using the residual method. **Hindlimb (HL, B):** No significant differences between the right and left hemispheres were observed (all Bonferroni-corrected *p* > 0.05). The ANOVA revealed a significant main effect of ROI (*F*(9,158) = 2.19, *p* = 0.025), but no main effect of between hemispheres (*F*(1,158) = 0.045, *p* = 0.83) or interaction (*F*(9,158) = 0.50, *p* = 0.87). **Forelimb (FL, C):** A trend toward reduced correlation in MOp in right hemispheres was observed (*t* = –4.21, uncorrected *p* = 0.0135, Bonferroni *p* = 0.1355), although not significant after correction. The ANOVA showed a significant main effect between hemispheres (*F*(1,80) = 4.88, *p* = 0.030), a trend for ROI (*F*(9,80) = 1.79, *p* = 0.083), and no significant interaction (*F*(9,80) = 0.87, *p* = 0.55). **Visual cortex (VC, D):** VISpm showed a trend toward decreased correlation in the right hemispheres (*t* = –2.31, *p* = 0.0380, Bonferroni *p* = 0.3804), with no other significant comparisons. The ANOVA revealed a significant main effect of ROI (*F*(9,256) = 2.66, *p* = 0.0057), but no significant main effect of between hemispheres (*F*(1,256) = 2.17, *p* = 0.14) or interaction (*F*(9,256) = 1.48, *p* = 0.16). **Barrel cortex (BC, E):** No statistically significant differences were found between the left and right hemispheres after correction (all Bonferroni *p* > 0.05), though SSp_bfd showed a trend (*t* = –2.13, *p* = 0.0663, Bonferroni *p* = 0.6634). The ANOVA indicated a significant main effect of ROI (*F*(9,224) = 2.08, *p* = 0.032), with no effect between hemispheres (*F*(1,224) = 0.73, *p* = 0.39) or interaction (*F*(9,224) = 0.85, *p* = 0.58). These results suggest that overall intrahemispheric GABAergic correlation patterns are largely stable across spontaneous and evoked conditions, with subtle region-specific modulations ,n=number of animals.

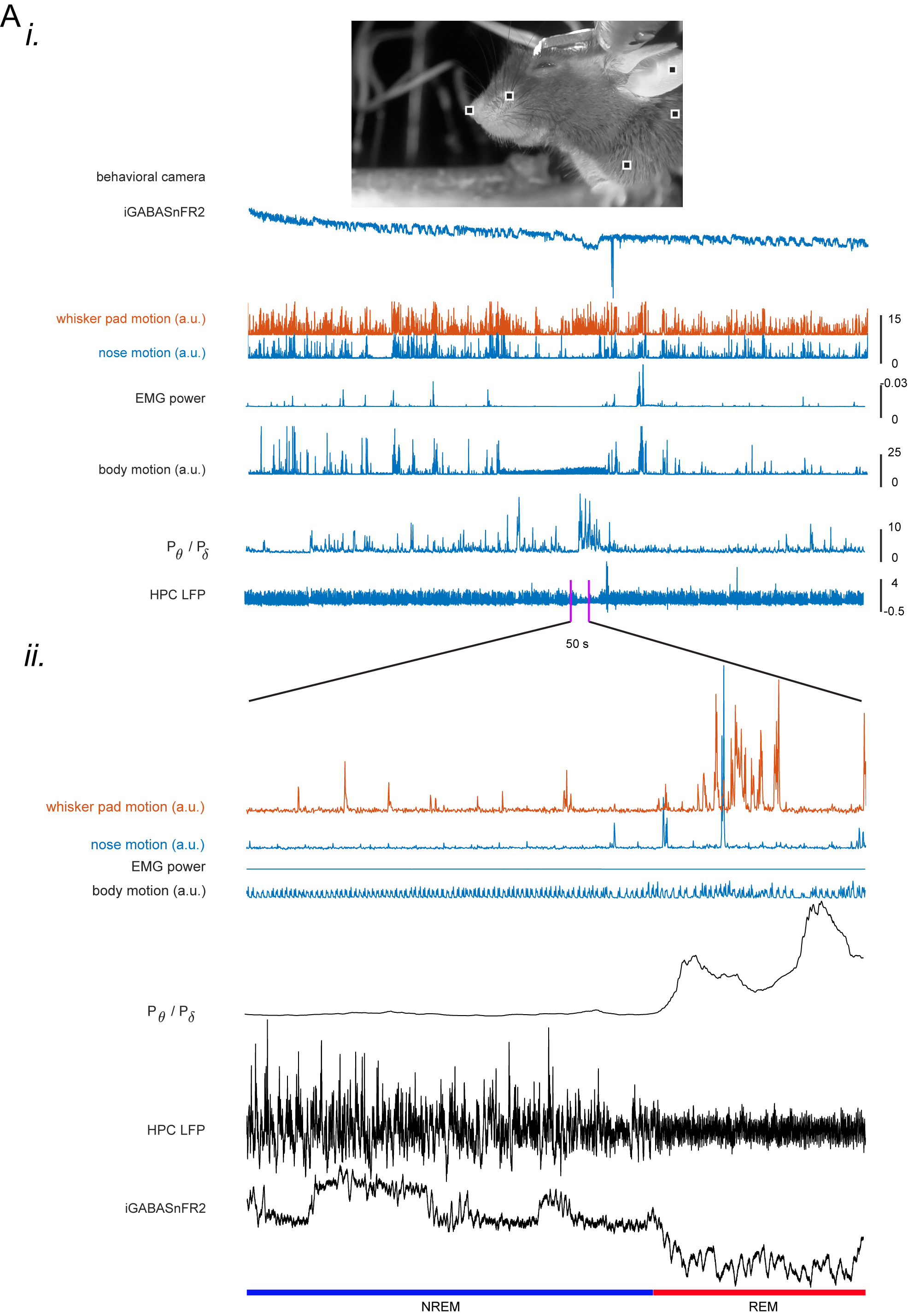

**Supplementary Figure 5. Motion signal, hippocampal LFP activity, and EMG power for sleep scoring in a head-fixed mouse.** Panel (i) presents a time-series analysis integrating motion and electrophysiological signals used for sleep scoring. The top schematic shows a spatial map of the mouse's head with rectangular regions of interest used to extract motion signals. The plots below display (from top to bottom): raw (non-preprocessed) iGABASnFR2 fluorescence , whisker pad and nose motion signals (z-scored) recorded at 25 Hz using a behavioral camera, EMG power indicating muscle activity, body motion (z-scored) representing gross movements, the theta-to-delta power ratio (Pθ/Pδ) derived from hippocampal local field potentials (LFPs) to differentiate NREM and REM sleep, and the raw hippocampal LFP signal reflecting underlying neural dynamics. Panel (ii) provides a magnified 50-second segment of the data shown in panel (i), offering a detailed view of whisker pad and nose motion, EMG power, body motion, theta-to-delta ratio, raw hippocampal LFP signal and raw (non-preprocessed) iGABASnFR2 signal. The highlighted segment captures transitions between NREM and REM sleep, demonstrating how motion and neural signals align with behavioral state changes.

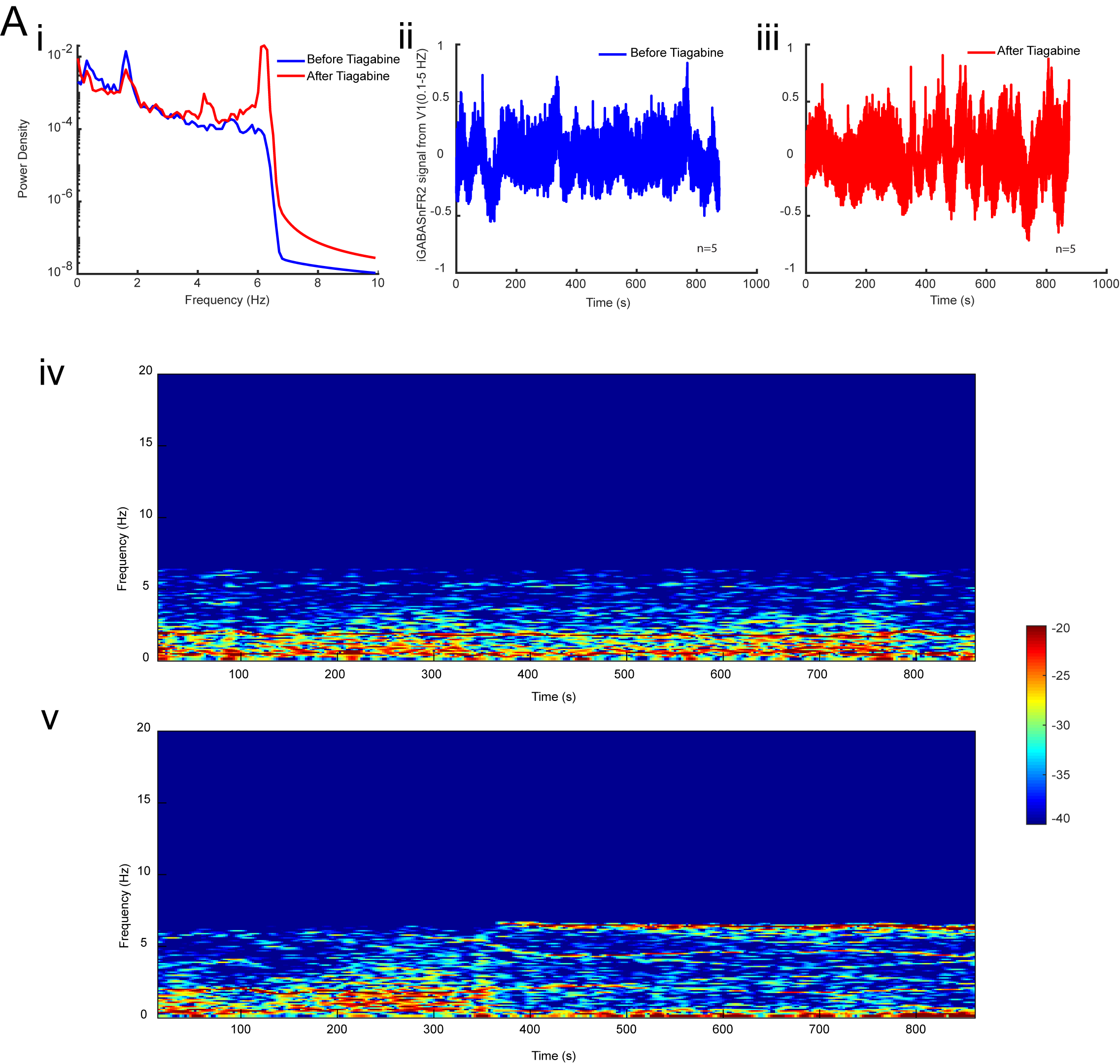
**Supplementary Figure 6. Tiagabine increases low-frequency GABAergic signal oscillations in the cortex. (A)** Power spectral and temporal dynamics of the iGABASnFR2 signal before and after Tiagabine administration. **(i)** Power spectral density (PSD) plot showing increased low-frequency (<1 Hz) oscillatory power following Tiagabine injection (red) compared to baseline (blue). **(ii, iii)** Z-scored time series of the global iGABASnFR2 signal before (ii, blue) and after (iii, red) Tiagabine administration. **(iv, v)** Time-frequency spectrograms of the iGABASnFR2 global signal before (iv) and after (v) Tiagabine injection, demonstrating a marked increase in low-frequency power post-injection, n=number of animals.
